# PLCγ1 promotes phase separation of the T cell signaling clusters

**DOI:** 10.1101/2020.06.30.179630

**Authors:** Longhui Zeng, Ivan Palaia, Anđela Šarić, Xiaolei Su

## Abstract

The T cell receptor (TCR) pathway receives, processes, and amplifies the signal from pathogenic antigens to the activation of T cells. Although major components in this pathway have been identified, the knowledge on how individual components cooperate to effectively transduce signals remains limited. Phase separation emerges as a biophysical principle in organizing signaling molecules into liquid-like condensates. Here we report that phospholipase PLCγ1 promotes phase separation of LAT, a key adaptor protein in the TCR pathway. PLCγ1 directly crosslinks LAT through its two SH2 domains. PLCγ1 also protects LAT from dephosphorylation by the phosphatase CD45 and promotes LAT-dependent ERK and SLP76 activation. Intriguingly, a non-monotonic effect of PLCγ1 on LAT clustering was discovered. Computer simulations, based on patchy particles, revealed how the cluster size is regulated by protein compositions. Together, these results define a critical function of PLCγ1 in promoting phase separation of the LAT complex and TCR signal transduction.

## Introduction

Mesoscale signaling clusters have been frequently observed in a variety of immune receptor pathways, including the T cell receptor (TCR) (Campi et al., 2005), B cell receptor (BCR) (Wang et al., 2017), Fc gamma receptor (Lin et al., 2016), engulfment receptor (Draper) (Williamson and Vale, 2018), T cell co-receptor CD28 (Yokosuka et al., 2008), and PD1 (Hui et al., 2017). These signaling clusters, ranging from several hundred nanometers to microns in size, share common features of complex composition, heterogeneous size, and dynamic assembly (Dustin and Groves, 2012). Because of these features, it remains a challenge to understand the mechanism and functional consequences of those clusters in transducing immune signaling.

The T cell microcluster represents a good example of this scenario. Following TCR activation, downstream signaling molecules self-organize into micron or submicron-sized clusters. These clusters are enriched of TCR, adaptor proteins LAT, Grb2, Gads, SLP76, Nck, and effectors ZAP70, Sos1, PLCγ1, CBL, and WASP (Balagopalan et al., 2015; Bunnell, 2010; Bunnell et al., 2002). Because the majority of early TCR signaling components reside in these microclusters, they are considered as a hub for transducing TCR signals (Balagopalan et al., 2015; Choudhuri and Dustin, 2010). Previous works showed that LAT, a transmembrane protein essential for TCR signal transduction (Zhang et al., 1998), serves as a scaffold to form a macromolecular signaling complex (Houtman et al., 2006; Zhang et al., 2000). Mathematical modeling suggested that multivalent protein-protein interactions play a critical role in forming the LAT complex (Nag et al., 2012; Nag et al., 2009). Using a supported lipid bilayer-based system, our previous work showed that LAT forms near micron-sized, membrane-embedded clusters through a mechanism of liquid-liquid phase separation (Su et al., 2016). The liquid-like LAT microclusters enrich kinase ZAP70 but exclude phosphatase CD45, thus promoting tyrosine phosphorylation. LAT microclusters also increase the downstream Ras activation (Huang et al., 2019), actin polymerization, and ERK activation (Su et al., 2016). These works revealed a critical role of LAT microclusters in promoting TCR signaling. However, the regulatory mechanism of LAT microclusters was not fully understood.

Following TCR activation, phospholipase C gamma 1 (PLCγ1) is recruited to the LAT microclusters, which further hydrolyzes PIP_2_ to generate IP_3_ and DAG, triggering calcium influx and PKC activation, respectively (Balagopalan et al., 2015; Courtney et al., 2018). PLCγ1 is a multi-domain lipase. Besides its catalytic core, PLCγ1 also contains two SH2 domains and one SH3 domain that serve structural or regulatory roles (Braiman et al., 2006; Manna et al., 2018). Its N-terminal SH2 domain (nSH2) binds specifically to phosphor-tyrosine (Y132) on LAT (Braiman et al., 2006) whereas its C-terminal SH2 domain (cSH2) is involved in releasing autoinhibition of PLCγ1 (Gresset et al., 2010; Hajicek et al., 2019). The SH3 domain of PLCγ1 directly interacts with the proline-rich motifs on Sos1 (Kim et al., 2000), a RasGEF which is also enriched in the LAT microclusters. The multiple binary interactions between PLCγ1 and other components in the LAT complex mediate a synergistic assembly of the LAT complex (Braiman et al., 2006; Hartgroves et al., 2003; Manna et al., 2018). However, because of the complex protein-protein interactions involved, the exact mechanism of how PLCγ1 regulates LAT microcluster formation remains elusive. This is mainly because traditional biochemical assays were performed in solution, which did not recapitulate two important features of cellular LAT microclusters: membrane association and giant size (up to micron).

We have recently developed a supported lipid bilayer-based system to reconstitute near-micron-sized LAT microclusters on synthetic membranes (Su et al., 2017). In the following study, using our biochemical reconstitution approach together with live cell microscopy and computer modeling, we delineated the role of PLCγ1 in regulating LAT microcluster formation. We found that the SH2 and SH3 domain of PLCγ1 can directly bridge LAT to Sos1 to form microclusters. PLCγ1 also protects LAT from CD45-dependent dephosphorylation. Therefore, PLCγ1 stabilizes LAT microclusters by both physical crosslinking and chemical protection. Moreover, we found that the PLCγ1 concentration influences the sizes of LAT microclusters in a non-monotonic way, pointing to a novel mechanism of the size control of liquid condensates. Together, these results expand the traditional view that PLCγ1 acts downstream of LAT; instead, PLCγ1 plays an active role in regulating the stability of LAT microclusters.

## Results

### PLCγ1 promotes LAT cluster formation *in vitro*

To determine how PLCγ1 regulates LAT microcluster formation, we implemented a supported lipid bilayer-based reconstitution assay that allows quantitative monitoring of microcluster assembly (Su et al., 2017). The cytoplasmic domain of LAT was purified, phosphorylated, and labeled with maleimide-Alexa488 on a C-terminal cysteine residue. It was then attached to the Ni2^+^-NTA functionalized supported lipid bilayer via a polyhistidine tag on the N-terminus. Total internal reflection fluorescence (TIRF) microscopy was used to visualize the formation of LAT clusters. As reported before (Su et al., 2016), LAT formed microclusters when Grb2 and Sos1 were added to the reaction mixture (Figure 1A and 1B). Intriguingly, when Grb2 was replaced with a fragment of PLCγ1 that contains the SH2 and SH3 domains, LAT still formed clusters, though with a higher number but smaller sizes, as compared to the Grb2-mediated cluster formation (Figure 1B). Furthermore, fluorescence recovery after photobleaching (FRAP) analysis revealed that PLCγ1-induced LAT clusters are less dynamic than Grb2-induced LAT clusters, as indicated by the lower recovery frequency and longer half recover time (Figure 1C). We also tested the full-length PLCγ1 and found that it induced LAT microcluster formation in a dose-dependent manner, which is similar to the fragment of PLCγ1 (Figure S1A and S1B). To understand if the ability to drive LAT cluster formation is a general feature of proteins containing the SH2 and SH3 domains, we replaced PLCγ1 with other LAT-associated proteins that play a role in TCR signal transduction. These include 1) Gads, an adaptor protein that binds LAT on overlapping sites with Grb2 (Zhang et al., 2000), 2) Vav1, a RhoGEF that closely associates with LAT (Sherman et al., 2016), and 3) Nck1, an adaptor protein that promotes actin polymerization (Wunderlich et al., 1999). We found that Gads, as reported before (Su et al., 2016), drives LAT microcluster formation whereas Vav1 or Nck1 did not (Figure S1C and S1D). Together, those data showed that PLCγ1, together with Sos1, can specifically induce LAT microcluster formation.

**Figure 1.**
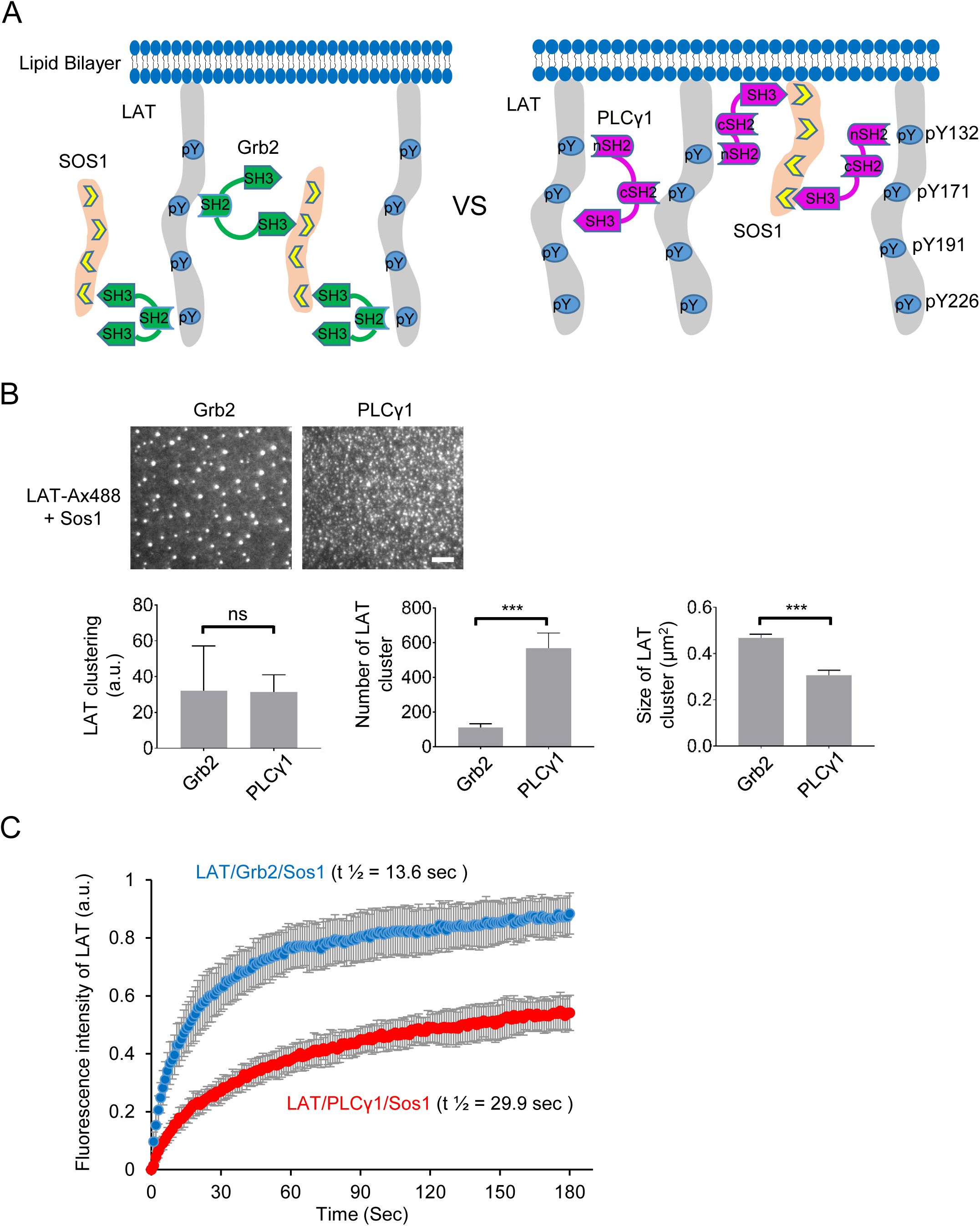
PLCγ1 promotes LAT cluster formation in vitro. (A) Schematics of the assay. (B) TOP: TIRF microscopy revealed that both Grb2 and PLCγ1 promote LAT microcluster formation. Alexa-488 labeled LAT at 300 molecules / μm^2^ was incubated with 125 nM Sos1 and 250 nM Grb2 or PLCγ1 for 0.5 hr before imaging. Scale bar: 5 μm. BOTTOM: Quantification of Grb2 or PLCγ1 driven-LAT microclusters. LAT clustering was quantified as normalized variance (Su et al., 2016). Shown are mean ± SD. N= 3 independent experiments. Unpaired two-tail t-test was used. ***: p<0.001; ns: not significant. (C) FRAP analysis revealed that PLCγ1 driven microclusters are less dynamic than Grb2-driven LAT microclusters. Shown are mean ± SD. N= 10 clusters.

### PLCγ1 crosslinks LAT through two SH2 domains

Next, we determined the mechanism by which PLCγ1 drives LAT microcluster formation. PLCγ1 contains an N-terminal SH2 domain (nSH2), a C-terminal SH2 domain (cSH2), and an SH3 domain. We produced PLCγ1 truncation mutants that lack either the nSH2, cSH2, or SH3 domain (Figure 2A). We found that mutants lacking either nSH2 or cSH2 lost the ability to drive LAT cluster formation whereas mutants lacking the SH3 domain still drove LAT cluster formation (Figure 2B and 2C). Because the SH3 domain interacts with the proline-rich motif on Sos1, the SH3-independent clustering suggested that Sos1 might be dispensable for PLCγ1-driven LAT cluster formation. Indeed, the nSH2-cSH2 fragment of PLCγ1 drove LAT cluster formation in the absence of Sos1 (Figure S2A and S2B). In this assay, PLCγ1 and mutants were used at 500 nM. We titrated PLCγ1 concentration and found that at a lower concentration of PLCγ1 (50 nM), Sos1 is still required for LAT microcluster formation (Figure S2C and S2D).

**Figure 2.**
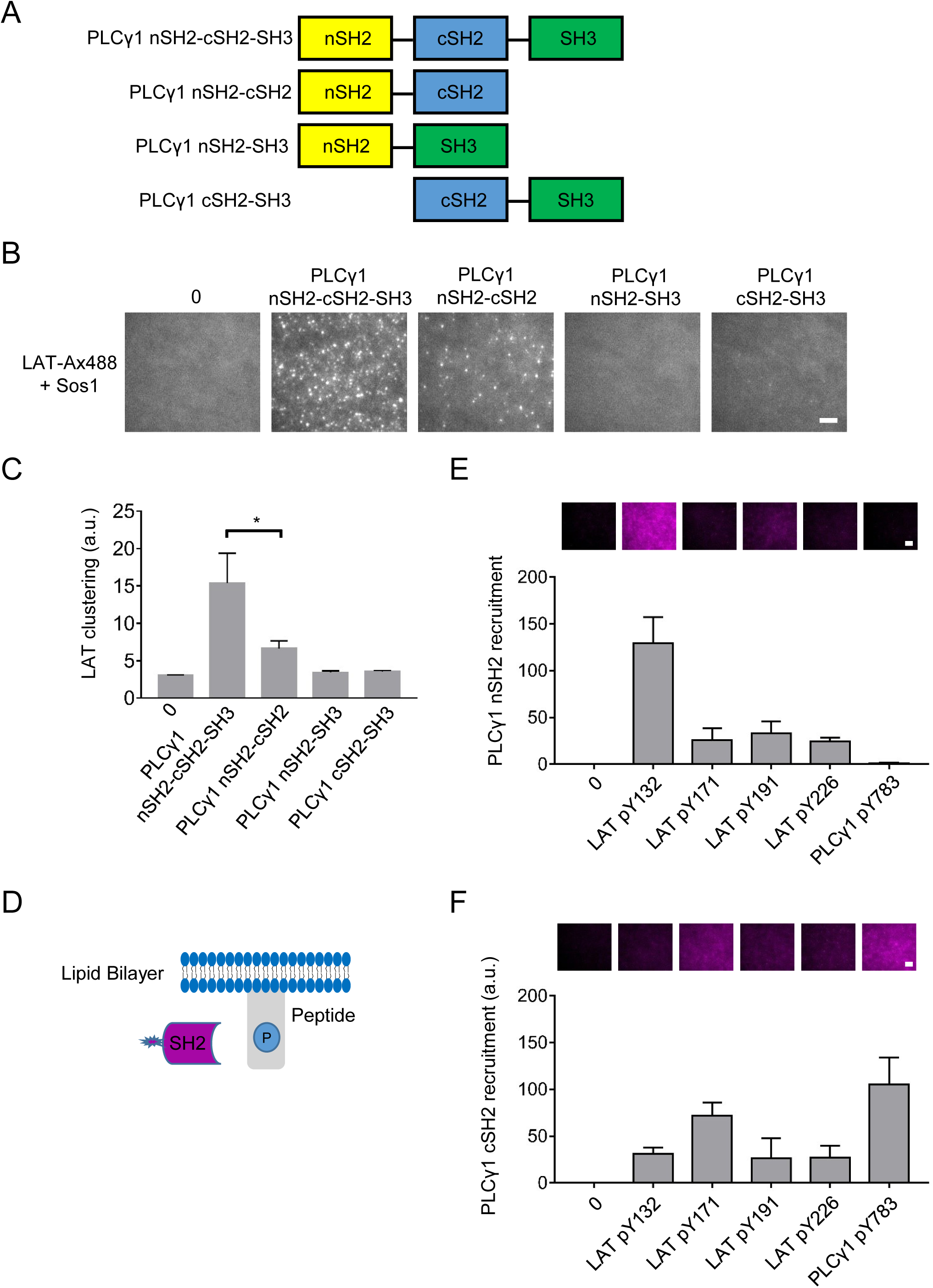
PLCγ1 crosslinks LAT by two SH2 domains. (A) Domains of the proteins used in the study. (B) TIRF microscopy revealed that both nSH2 and cSH2 domains are required for PLCγ1 driven-LAT microcluster formation. SH3 domain promotes cluster formation. Alexa-488 labeled LAT at 300 molecules / μm^2^ was incubated with 300 nM Sos1 and 50 nM PLCγ1 for 0.5 hr before imaging. Scale bar: 5 μm. (C) Quantification of PLCγ1-driven LAT microclusters. Shown are mean ± SD. N= 3 independent experiments. Unpaired two-tail t-test was used. *: p<0.05; (D) Schematics of the assay of testing SH2 domain binding sites. (E) PLCγ1 nSH2 binds LAT Y132. Phospho-peptides were synthesized, biotinylated at the N-terminus, and attached to the biotin-functionized supported lipid bilayers by streptavidin. The SH2 domain were labeled with fluorescent dye (Maleimide Ax647), and incubated with the individual phosphor-peptides. The membrane-associated SH2 domain was measured by TIRF microscopy. Scale bar: 5 μm. (F) PLCγ1 cSH2 binds LAT Y171. Same settings as in (E).

Next, we determined the binding sites on LAT that interact with the SH2 domain of PLCγ1. The four C-terminal tyrosines on LAT are necessary and sufficient to transduce the TCR signaling (Zhu et al., 2003). Synthesized peptides containing each one of these four phosphotyrosines of LAT were attached to the supported lipid bilayer. Recombinant nSH2 or cSH2 domain of PLCγ1 was purified, labeled with CoA-647 on an N-terminal ybbR tag, and incubated with individual phospho-peptides on the membrane (Figure 2D). The binding of the SH2 domain to phosphopeptide on the membrane was revealed by TIRF microscopy. We found that the nSH2 domain strongly interacted with LAT Y132 (Figure 2E), which is consistent with previous reports (Zhang et al., 2000). The cSH2 domain robustly bound to LAT Y171, though the binding was slightly lower than PLCγ1 Y783 (Figure 2F), the previously reported site that interacts with the cSH2 domain in cis (Hajicek et al., 2013; Poulin et al., 2005). Together, those data suggested a model in which PLCγ1 crosslinks LAT through two interaction pairs: nSH2 with Y132, and cSH2 with Y171.

### PLCγ1 cooperates with Grb2 to regulate LAT clustering

Next, we investigated how PLCγ1 cooperates with Grb2 and Sos1 to induce LAT microcluster formation. Because cluster formation depends on the concentration of individual components, we adopted concentrations of proteins that were measured in T cells (Nag et al., 2009; Voisinne et al., 2019). We incubated LAT at a density of 300 molecules / μm^2^ with 3000 nM Grb2 and 300 nM Sos1. PLCγ1 was additionally included in the clustering assay. Surprisingly, we found that PLCγ1, at 50 nM (concentration in T cells), could significantly increase LAT cluster formation and recruitment of Sos1 to the membrane. This clustering-promoting effect is robust even at a concentration as low as 5 nM PLCγ1 (Figure 3A and 3B). Notably, this concentration (5 nM) is orders of magnitude lower than Grb2 (3000 nM), the other SH2-SH3 adaptor in the system. Furthermore, FRAP analysis revealed that PLCγ1 significantly reduces the exchange of LAT molecules between inside and outside of the clusters (Figure S3), supporting the idea that PLCγ1 serves as an orthogonal crosslinker to Grb2, to stabilize LAT clusters. To understand how PLCγ1 affects the kinetics of LAT cluster formation, we performed time lapse imaging. LAT was attached to the membrane, Grb2/Sos1 was added into the system at time 0. We observed cluster formation as usual (Figure 3C and Movie S1). Intriguingly, when PLCγ1 was additionally included in the system, clustering was significantly enhanced from the very beginning (Figure 3C and Movie S2). Together, these data suggest that, PLCγ1, by serving as a crosslinker, could dramatically increase LAT microcluster formation.

**Figure 3.**
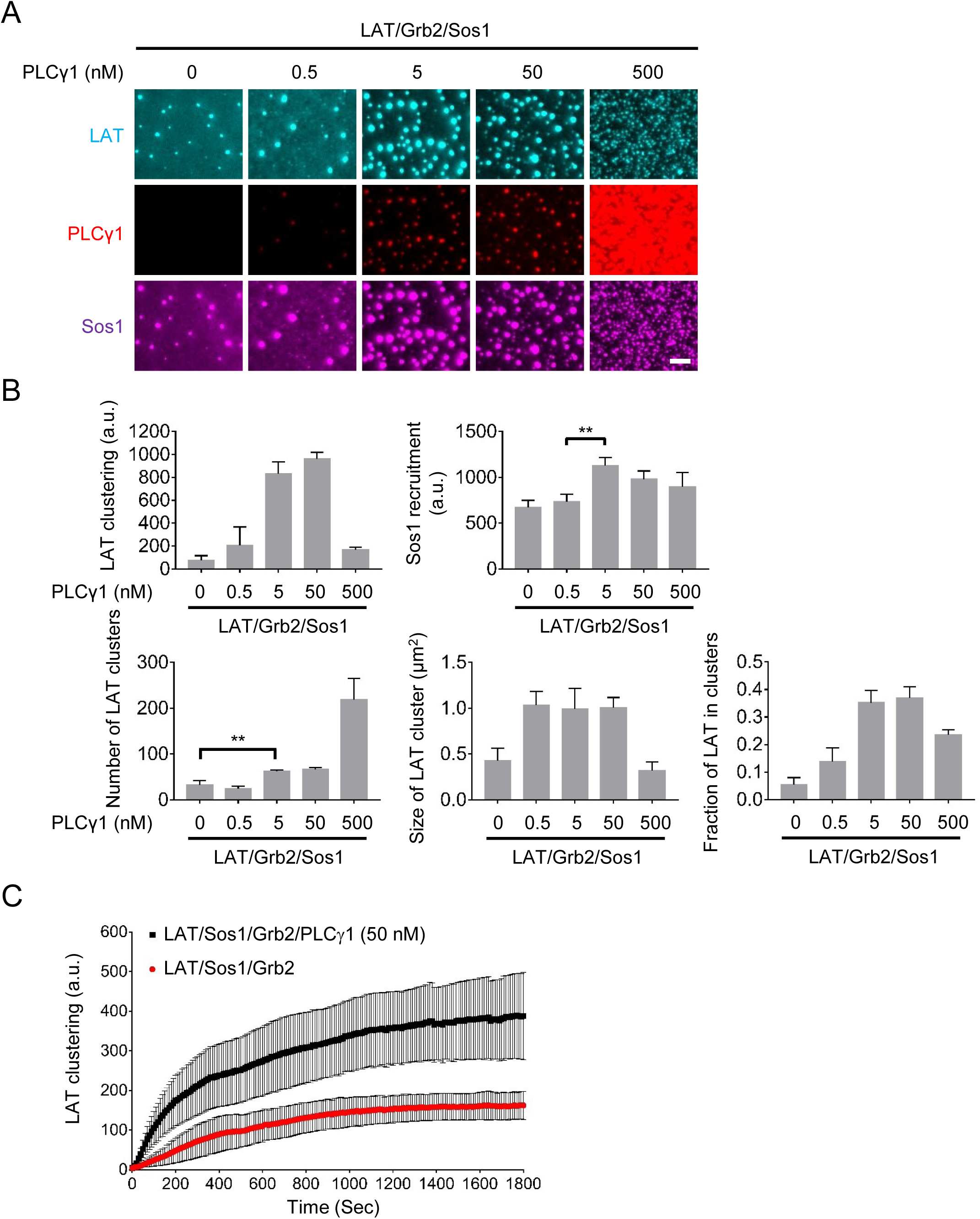
PLCγ1 cooperates with Grb2 to regulate LAT microcluster formation. (A) TIRF microscopy revealed that PLCγ1 regulates LAT microcluster formation in a non-monotonic manner. Physiologically relevant concentrations of proteins were used in the assay: LAT at 300 molecules / μm^2^, Grb2 at 3 μM, Sos1 at 0.3 μM, and PLCγ1 at 50 nM. LAT was labeled with Alexa-488, PLCγ1 was labeled with DY547, and Sos1 was labeled with Alexa-647. Scale bar: 5 μm. (B) Quantification of LAT clustering, membrane recruitment of Sos1. Shown are mean ± SD. N= 3 independent experiments. Unpaired two-tail t-test was used. **: p<0.01. (C) PLCγ1 accelerates LAT cluster formation. TIRF microscopy revealed the time course of LAT microcluster formation in the presence or absence of PLCγ1. LAT-Alexa488, at 1000 molecules / μm^2^ was incubated with 1000 nM Grb2 and 500 nM Sos1, and/or 50 nM PLCγ1 at time 0. Shown are mean ± SEM. N= 3 independent experiments.

### Computer Simulations of PLCγ1-mediated LAT clustering

Intriguingly, a non-monotonic effect of PLCγ1 on LAT clustering was revealed. PLCγ1, at low concentrations, promotes LAT clustering and increases cluster sizes but this effect is diminished at high concentrations (Figure 3A and 3B). This points to the fact that PLCγ1 not only promotes LAT clustering, but also regulates the cluster size. To understand the physical mechanism underlying this size regulation, we developed a minimal coarse-grained computer model in which the proteins are described as spherical particles decorated with patches that represent binding domains and simulated via molecular dynamics. The model, described quantitatively in Methods and in Supplemental Information (SI), consists of 4 types of two-dimensional particles, representing LAT, PLCγ1, Sos1 and Grb2; these can bind mutually, respecting biochemical valence and bond specificity (Figure 4A). In simulations, the pool of such two-dimensional particles readily aggregates into clusters that grow either via addition of individual proteins or via merging with other clusters until they reach steady-state sizes. The average cluster size in simulations exhibits a non-monotonic dependence on the concentration of PLCγ1 (Figure 4B, and Movie S3-S8), indeed recapitulating what was observed in experiments (Figure 3B). The non-monotonic effect was robustly revealed in a wide range of LAT densities with cluster size measured either by the number of LAT or by the total number of four proteins (Figure S4A). To understand the origin of this non-monotonic phenomenon, we computed, at a given time, the number of possible bonds between free binding sites that can make whichever two clusters merge into a bigger one; this parameter, that we named coalescence likelihood, is a measure of how easy it is for the clusters to merge at a certain PLCγ1 concentration. We found that the coalescence likelihood closely captures the non-monotonicity of the cluster size (Figure 4B). A breakdown of the coalescence likelihood by bond type (Figure 4C), together with an analysis of the average coordination per molecule (Section SI 3), showed the following.

**Figure 4.**
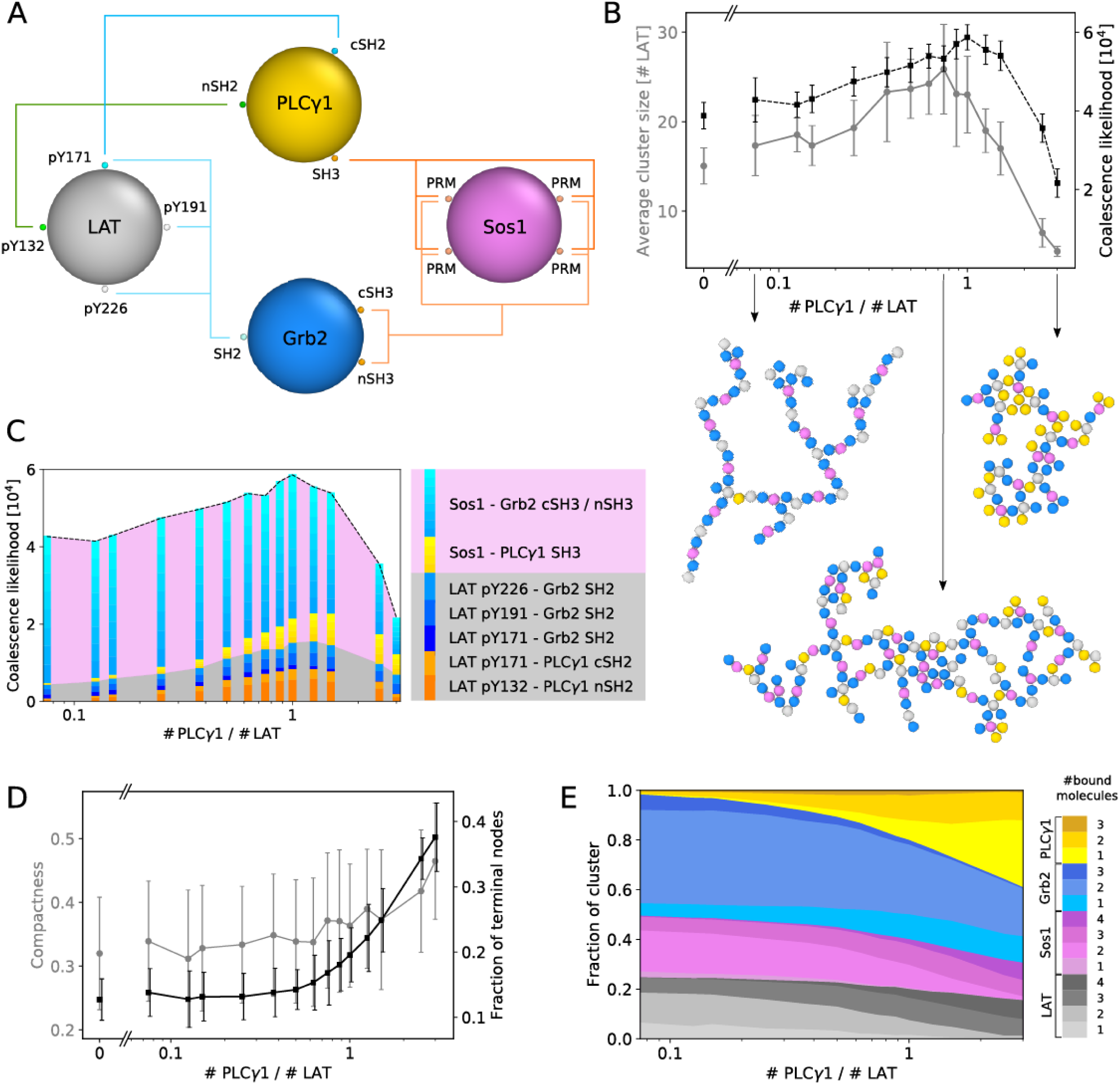
A coarse-grained model explains how PLCγ1 non-monotonically regulates LAT clustering. (A) Sketch of the model in which the proteins are represented as two-dimensional particles decorated by interaction patches. All bonds possible in the system, based on biochemical data, are illustrated with colored lines. (B) Top panel: the average cluster size displays non-monotonic dependence on the PLCγ1 concentration (gray circle). This behavior is well captured by the likelihood for clusters coalescence (black squares). Error bars represent statistical errors on the average size over different realization of the simulation. Bottom panel: snapshots of typical clusters in simulations, for relative PLCγ1:LAT concentrations of 0.075, 0.75, and 3 (these clusters contain respectively 19, 30 and 10 LAT molecules, and, with reference to panel D, their compactness is 0.23, 0.39, and 0.48). (C) Breakdown of the coalescence likelihood per type of possible bond. The gray and pink areas represent available bonds involving a LAT or a Sos1 molecule, respectively; blue and yellow-orange bars represent bonds involving Grb2 and PLCγ1, respectively. (D) Compactness (gray circles, see Methods) and fraction of terminal nodes (black squares), as a function of PLCγ1 concentration. (E) Fraction of LAT, PLCγ1, Sos1, and Grb2 molecules per cluster, as a function of PLCγ1 concentration, shaded according to the number of other molecules they are bound to.

At low concentrations, PLCγ1 particles provide a binding site for the otherwise unbound pY132 in LAT particles and increase binding of the pY171 site and of all Sos1 sites, through an SH3-PRM (proline-rich motif) bond. The availability of these additional binding sites increases the coalescence likelihood (yellow and orange bars in Figure 4C); at the same time, it increases the average coordination (number of sites that interact with other molecules) of both LAT and Sos1 particles, making clusters more connected (Figure 4E and S4B). At high concentrations, though, PLCγ1 ends up saturating PRMs on Sos1 and, more slowly, phosphotyrosines on LAT particles, with the help of Grb2 (Figure S4B). As this occurs, free binding sites become rare and the coalescence likelihood decreases drastically. Clusters are still more connected and compact than without PLCγ1, but they are smaller: indeed, Sos1 and LAT are almost fully bound and it is unlikely for new bonds to form upon random collisions (pink and gray bands in Figure 4C restrict by more than half, Movie S8).

A further analysis on the compactness of clusters was performed by computing their inertia tensor, as well as by graph-theoretical means (see Methods and Figure S4B). The compactness index showed that clusters become monotonically more compact as PLCγ1 concentration increases (Figure 4D). Nonetheless, at high concentrations, clusters exhibit a large amount of ‘terminal nodes’, i.e. particles bound to one other particle only. This is due to the overwhelming amount of PLCγ1 particles, which tend to cap any binding site available to them, competing with themselves and with Grb2 for LAT and Sos1 sites. At the same time, as expected, LAT and Sos1 molecules become more and more bound (Figure 4E and S4B). This suggests that the stabilizing effect of PLCγ1 observed in FRAP experiments is due to increased compactness of clusters and increased complexity of the LAT network therein. In short, PLCγ1 concentration emerges as a possible regulator both of cluster size and of cluster stability.

### PLCγ1 promotes LAT clustering in T cells

We then investigated how PLCγ1 regulates LAT cluster formation in T cells. A LAT-mCherry construct was introduced into the wild-type or PLCγ1 null Jurkat T cells by lentiviral transduction. Those T cells were activated by cover glass-coated TCR-activating antibody OKT3. The formation of LAT microclusters was monitored by TIRF microscopy. We found that LAT clustering in PLCγ1 null cells was significantly reduced as compared to the wild-type cells (Figure 5A). This suggested a positive role of PLCγ1 in promoting LAT clusters, which is consistent with forehead in vitro results. To understand how the SH2 and SH3 domain of PLCγ1 contribute to LAT clustering, we reconstituted PLCγ1 null cells with the full-length PLCγ1 or PLCγ1 lacking the nSH2, or SH3 domain. These cells also express LAT-mCherry, as an indicator for LAT clusters. Unfortunately, PLCγ1 null cells reconstituted with PLCγ1 lacking the cSH2 domain do not grow, potentially because of the hyperactivity and toxicity resulting from the deletion of the inhibitory cSH2 domain. We found that LAT microcluster formation was significantly higher in the wild-type cells, as compared to the *Δ*nSH2 or *Δ*SH3 cells (Figure 5B).

**Figure 5.**
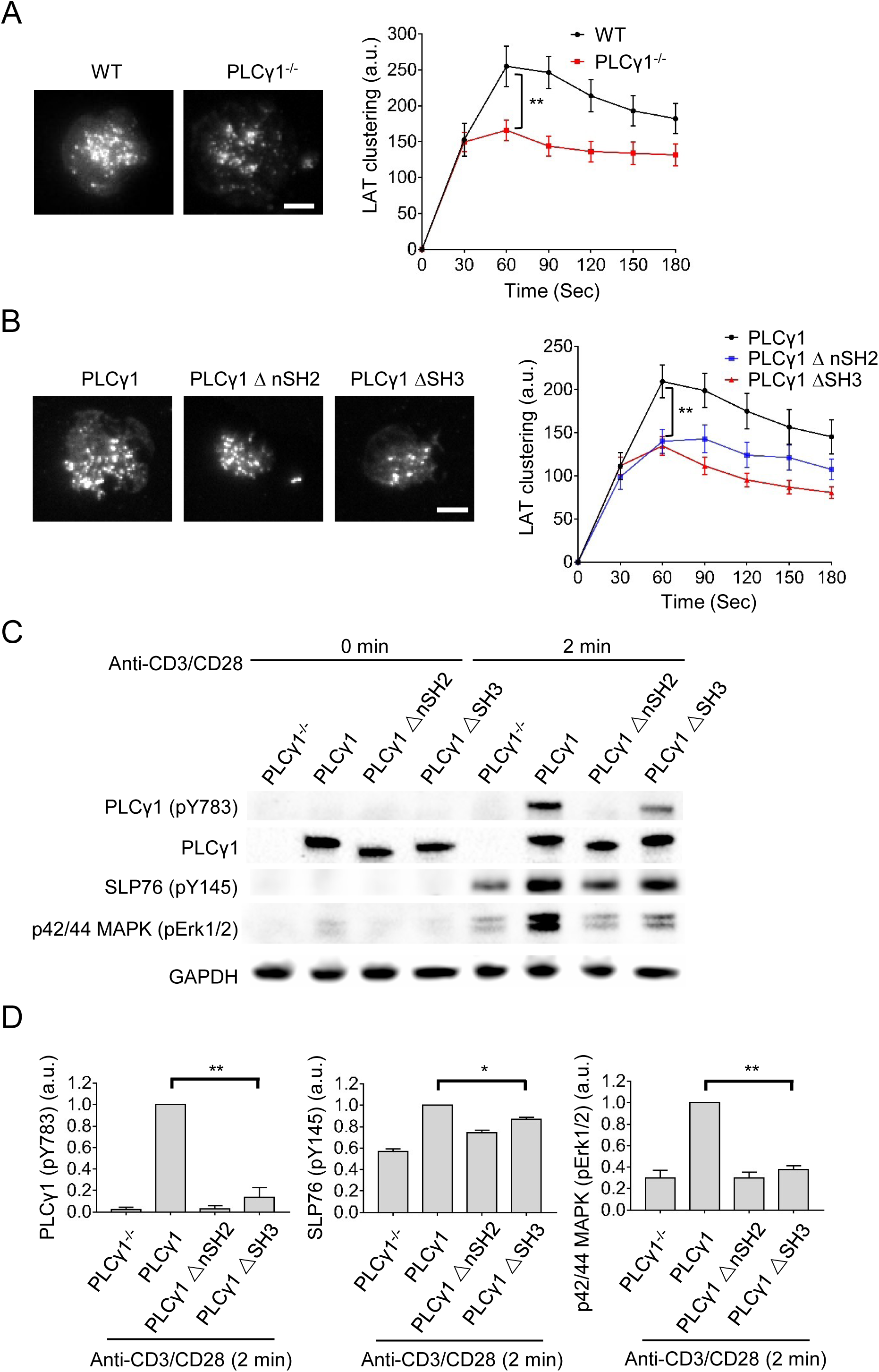
PLCγ1 promotes LAT clustering, SLP76 and ERK activation in Jurkat T cells. (A) Diminished LAT microcluster formation in PLCγ1 null cells. Wild-type or PLCγ1 null Jurkat T cells expressing LAT-mCherry were plated on OKT3-coated cover glass. LAT microcluster formation was revealed by TIRF microscopy. Images showed clustering 90s after cell landing on the glass. Scale bar: 5 μm. Shown are mean ± SEM. N= 25 to 26 cells. Unpaired two-tail t-test was used. **: p<0.01. (B) The nSH2 and SH3 domain of PLCγ1 promotes LAT cluster formation. PLCγ1 null Jurkat T cells expressing LAT-mCherry were reconstituted with the wild-type, *Δ*nSH2, or *Δ*SH3 PLCγ1. Those cells were plated on OKT3-coated cover glass. Images showed clustering 90s after cell landing on the glass. LAT microcluster formation was revealed by TIRF microscopy. Scale bar: 5 μm. Shown are mean ± SEM. N= 22 to 30 cells. Unpaired two-tail t-test was used. **: p<0.01. (C) Immunoblot analysis of LAT-null Jurkat T cells reconstituted with the wild-type, *Δ*nSH2, or *Δ*SH3 PLCγ1. Cells were stimulated with 2 μg/mL anti-CD3 and anti-CD28 antibodies for 2 min, lysed, and applied for western blot analysis. (D) Quantification of the level of indicated proteins, after normalized to the expression level of GAPDH. Shown are mean ± SD. *N*= 3 independent experiments. Unpaired two-tail t-test was used. *p < 0.05, **p < 0.01.

Because LAT clustering activates downstream signaling pathway including SLP76 and MAPK, we decided to determine if PLCγ1 affects these pathways. Jurkat T cells were activated by anti-CD3 and anti-CD28 antibodies before being harvested for immunoblot analysis. Indeed, as compared to cells expressing the wild-type PLCγ1, cells expressing PLCγ1*Δ*nSH2 or PLCγ1*Δ*SH3 displayed defects in SLP76 activation (as indicated by SLP76 pY145) and MAPK activation (as indicated by pERK1/2) (Figure 5C and 5D). We will discuss the possible pathway leading to these defects in Discussion. Together, these data suggest that the structural domains of PLCγ1 promote LAT clustering and LAT downstream pathways in T cells.

### PLCγ1 protects LAT from dephosphorylation

The formation of LAT microcluster is phosphorylation-dependent. Antigen engagement with TCR triggers the phosphorylation of LAT by the kinase ZAP70, which is antagonized by phosphatases (Su et al., 2016). LAT, once phosphorylated, can recruit SH2-containing proteins to form microclusters. Therefore, PLCγ1 has traditionally been viewed as a “passenger” that is passively recruited to phosphorylated LAT. Interestingly, we found that PLCγ1 also regulates the phosphorylation of LAT. As compared to the wild-type cells, PLCγ1 null cells had significantly lower phosphorylation on Y132, but not on Y191 (Figure 6A), a binding site for Grb2 and Gads (Zhang et al., 2000). Deleting Grb2 or Gads, the other major SH2 domain-containing proteins in the LAT complex, did not alter the phosphorylation of Y132 (Figure 6A). This is consistent with the fact that Y132 only binds PLCγ1 but not Grb2 or Gads. Intriguingly, overexpressing a fragment of PLCγ1 that contains the SH2 and SH3 domains significantly increased phosphorylation on Y132 (Figure 6A), suggesting PLCγ1 as a two-way (both up and down) regulator of Y132 phosphorylation. Because PLCγ1 is the only identified binding partner of LAT Y132, we hypothesized that the SH2 domain of PLCγ1 binds to LAT Y132, protecting it from being dephosphorylated. Supporting that, when the cellular phosphatase activity was inhibited by vanadate, the difference in phosphorylation level of Y132 between the wild-type and PLCγ1 null cells was abolished (Figure 6B). To directly test the hypothesis that PLCγ1 protects LAT Y132 from dephosphorylation, we set up an in vitro dephosphorylation assay. LAT-Grb2-Sos1 microclusters were assembled in the presence or absence of PLCγ1. Then CD45, the most abundant phosphatase on T cell membranes, was added to dephosphorylate LAT (Figure 6C). Intriguingly, PLCγ1 significantly suppressed the dephosphorylation of LAT Y132 (Figure 6D). Together, these data suggest that PLCγ1 specifically stabilizes the phosphorylation on LAT Y132 by protecting it from being dephosphorylated by CD45.

**Figure 6.**
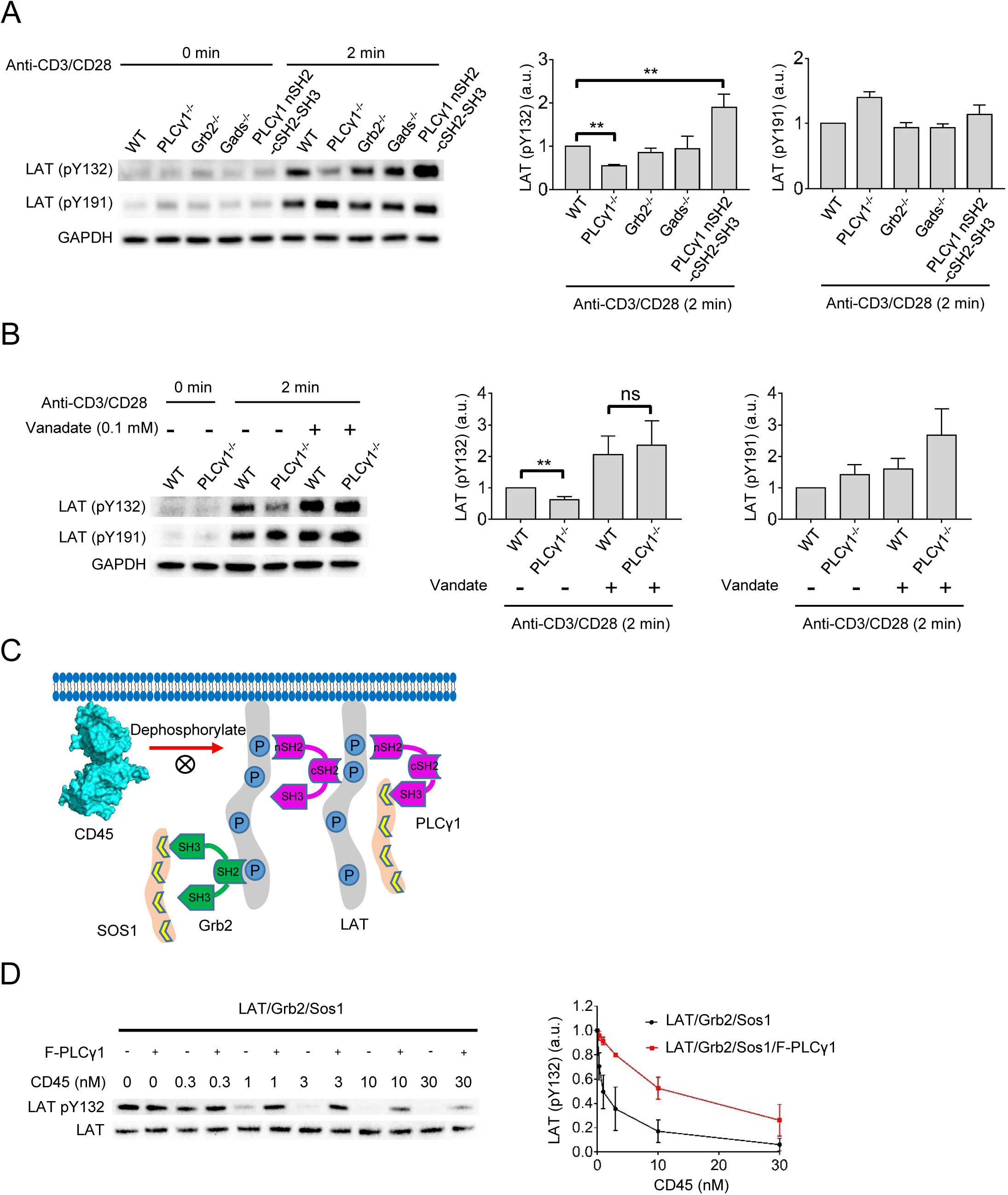
PLCγ1 protects LAT from dephosphorylation by CD45. (A) Reduced phosphorylation at LAT Y132 in PLCγ1 null cells. Cells as indicated were stimulated with 2 μg/mL anti-CD3 and anti-CD28 antibodies for 2 min, lysed, and applied for western blot analysis. The level of indicated proteins, after normalized to the level of GAPDH, was quantified. Shown are mean ± SD. *N*= 3 independent experiments. Unpaired two-tail t-test was used. **p < 0.01. (B) PLCγ1 prevents LAT Y132 from being dephosphorylated. Cells as indicated were pretreated with 0.1 mM vanadate (pan phosphatase inhibitor) before being stimulated with 2 μg/mL anti-CD3 and anti-CD28 antibodies for 2 min, lysed, and applied for western blot analysis. The level of indicated proteins, after normalized to the level of GAPDH, was quantified. Shown are mean ± SD. *N*= 3 independent experiments. Unpaired two-tail t-test was used. **p < 0.01; ns: not significant. (C) Schematics of the in intro dephosphorylation assay. (D) PLCγ1 prevents LAT Y132 from being dephosphorylated by CD45 in vitro. pLAT, at 1000 molecules / μm^2^, was incubated with 1 μM Grb2, 0.5 μM Sos1, and / or 100 nM full-length PLCγ1 for 0.5 hr. CD45 was then added to dephosphorylate pLAT for 5 min. The reaction was terminated by adding SDS-PAGE loading buffer with 2 mM vanadate. The level of phosphorylated LAT, after normalized to total LAT, was quantified. Shown are mean ± SD. N= 3 independent experiments.

## Discussion

PLCγ1 was traditionally considered as an enzyme that acts downstream of LAT. Following TCR activation, PLCγ1 is recruited to LAT microclusters and activated by Itk1-triggered phosphorylation. Activated PLCγ1 hydrolyzes PIP_2_ to generate IP_3_ and DAG, which triggers downstream calcium and PKC pathway, respectively. Here we reported that PLCγ1 can also promote LAT cluster formation. This is achieved by two ways: PLCγ1 crosslinks LAT directly or indirectly through Sos1; and PLCγ1 protects LAT from being dephosphorylated by CD45. Therefore, PLCγ1 and LAT reciprocally regulate each other in a positive manner.

The cSH2 domain of PLCγ1 is well-known for its role in regulating the enzymatic activity of PLCγ1 (Gresset et al., 2010; Hajicek et al., 2013). Combining our new results showing that cSH2 promotes LAT clustering, we propose the following refined model explaining the recruitment and activation of PLCγ1 during T cell activation (Figure S6A): following TCR engagement with antigens, LAT is phosphorylated at Y171, Y132 and other sites. These two sites interact with the cSH2 and nSH2 domain of PLCγ1, respectively, to recruit PLCγ1 from the cytosol to the membrane. Then together with Grb2, Sos1, and other LAT binding partners, PLCγ1 promotes LAT cluster formation. This will further recruit Itk1, which was shown to bind the LAT complex (Bunnell et al., 2000; Ching et al., 2000), to the LAT microclusters. Itk1 phosphorylates PLCγ1 on Y783 (Perez-Villar and Kanner, 1999). Phosphorylated Y783 interacts with the cSH2 domain intramolecularly to release the autoinhibition of PLCγ1 and activate the lipase activity. The cSH2 domain switches from LAT-binding to PLCγ1-binding, which reduces the avidity of PLCγ1 to the LAT complex. Consequently, PLCγ1 disassociates from the LAT clusters, either translocates to the TCR clusters (Cruz-Orcutt et al., 2014) or moves back to the cytosol.

Our domain truncation analysis of PLCγ1 in T cells revealed that the SH2 and SH3 domain could contribute to SLP76 and ERK activation. This could be explained by two non-exclusive mechanisms: 1) the SH3 domain of PLCγ1 directly binds Sos1 (Kim et al., 2000) and recruits Sos1 to the membrane, which facilitates Ras and ERK activation; similarly, the SH3 domain directly binds and recruits SLP76 to the membrane (Yablonski et al., 2001) in preparation for being phosphorylated by ZAP70; 2) the SH2 and SH3 domains promote LAT cluster formation, which further enhances the recruitment of Grb2 and Gads to LAT. Grb2 and Gads are constitutive binding partners for Sos1 and SLP76, respectively. Therefore, the SH2 and SH3 domain of PLCγ1 can contribute to downstream signaling both directly or indirectly through LAT.

Through both biochemical reconstitution and computational approaches, we revealed a non-monotonic mechanism of regulating LAT clustering by PLCγ1. Compositional control emerges as an important mechanism for regulating the physical feature and chemical activities of liquid-like condensates (Banani et al., 2016; Ditlev et al., 2019; Riback et al., 2020). The concentration of PLCγ1 may have been tuned to control cluster size and stability, with the purpose of transducing signaling to the physiological needs. Interestingly, PLCγ1 is upregulated in acute myeloid leukemia (Mahmud et al., 2017), colorectal carcinoma (Noh et al., 1994), and squamous cell carcinoma (Xie et al., 2010). It remains as an interesting question whether PLCγ1 mis-regulates signaling cluster formation in these pathological conditions.

PLCγ1 is involved in a variety of membrane receptor signaling pathways. The natural killer cell and mast cell receptor (FcεR1) pathway share almost the same machinery of LAT clustering with the T cell. We reasoned that PLCγ1 could play a similar role in promoting receptor signaling by enhancing LAT clusters in natural killer and mast cells. In the pathways outside immune responses, such as EGFR, FGFR or HER2, LAT is absent. However, similar scaffold proteins are present that contain multivalent interaction sites (e.g. EGFR and Shc) that can interact with PLCγ1 and Grb2-Sos1. The cooperation between PLCγ1 and Grb2 in promoting receptor clustering, as revealed in this work, could serve as a general mechanism for regulating membrane receptor signaling.

## Acknowledgements

We thank Christian Vanhille Campos and Johannes Krausser for providing basis for the simulation code and discussion on this project. X.S. has received support from the American Cancer Society Institutional Research Grant, the Charles H. Hood Foundation Child Health Research Awards, the Andrew McDonough B+ Foundation Research Grant, the Gilead Sciences Research Scholars Program in Hematology/Oncology, and the Rally Foundation a Collaborative Pediatric Cancer Research Awards Program. I.P. and A.Š. received funding from the European Research Council (StG 802960) and Royal Society.

## Author Contributions

Z.L. and X.S. conceived the projects and wrote the manuscripts with the contributions from I.P. and A.Š. Z.L. performed wet experiments and analyzed the data. I.P. and A.Š. conceived the computer model. I.P. performed computer simulations and analyzed the data. X.S. and A.Š. supervised the project.

## Materials and Methods

### Key resources table

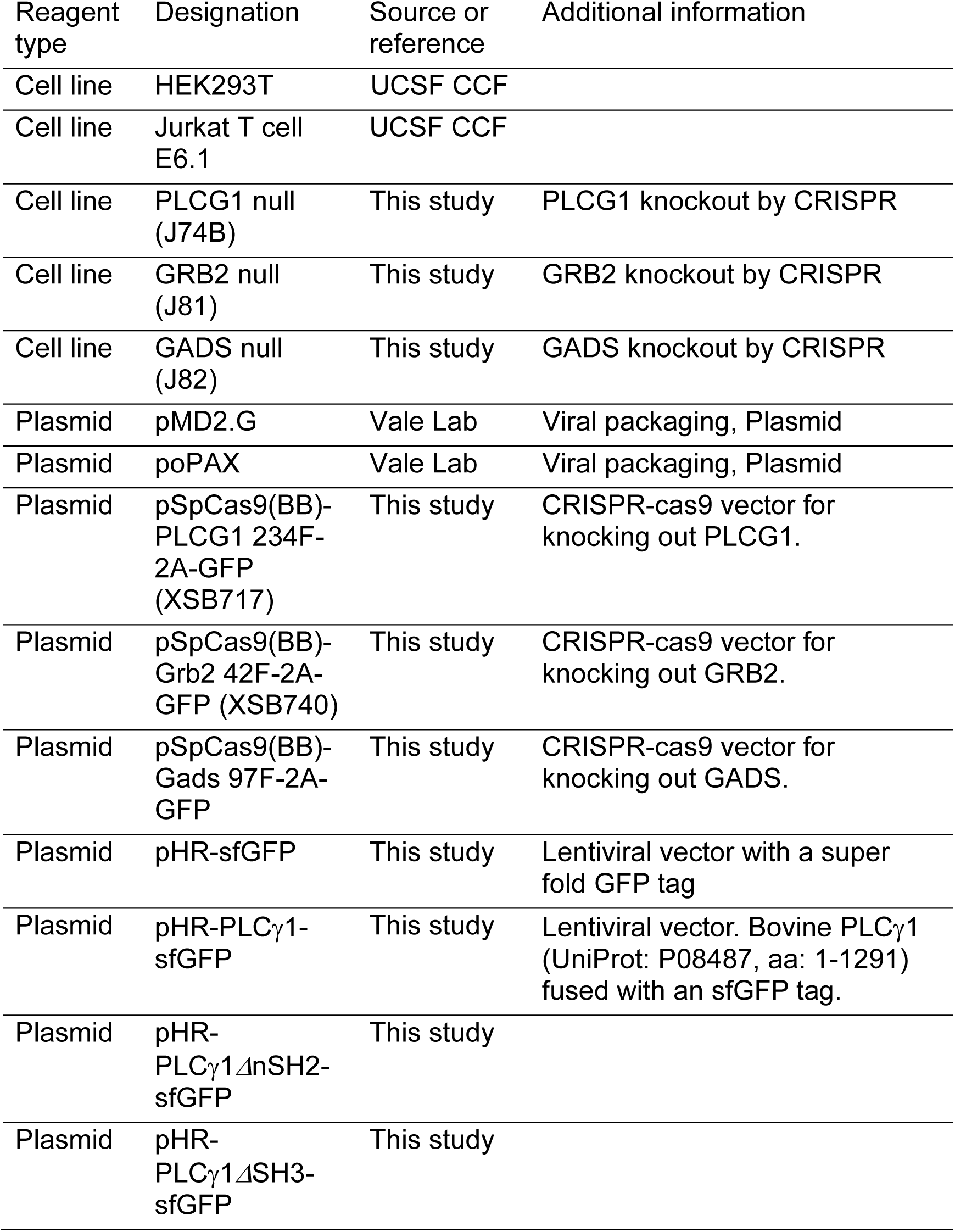
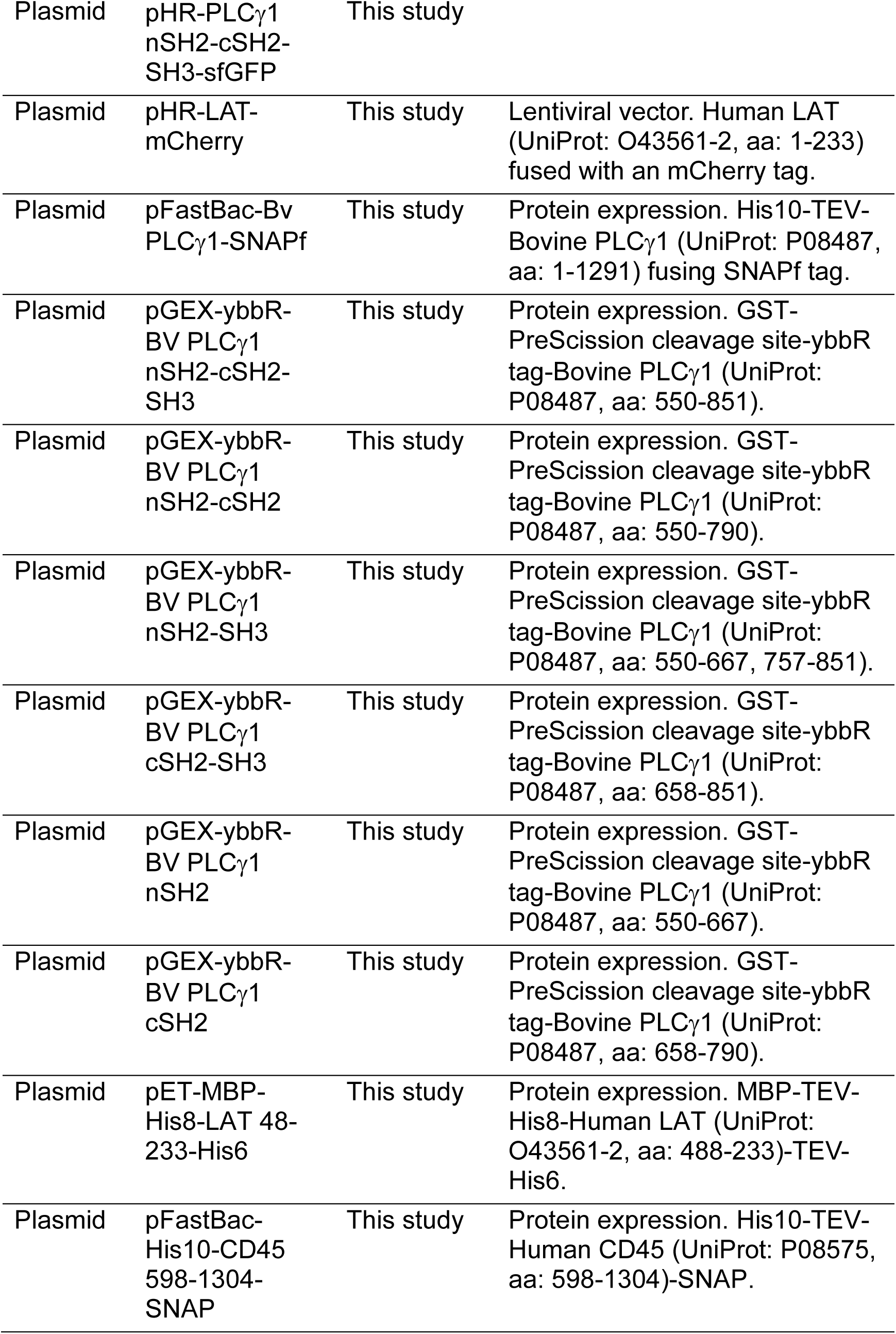
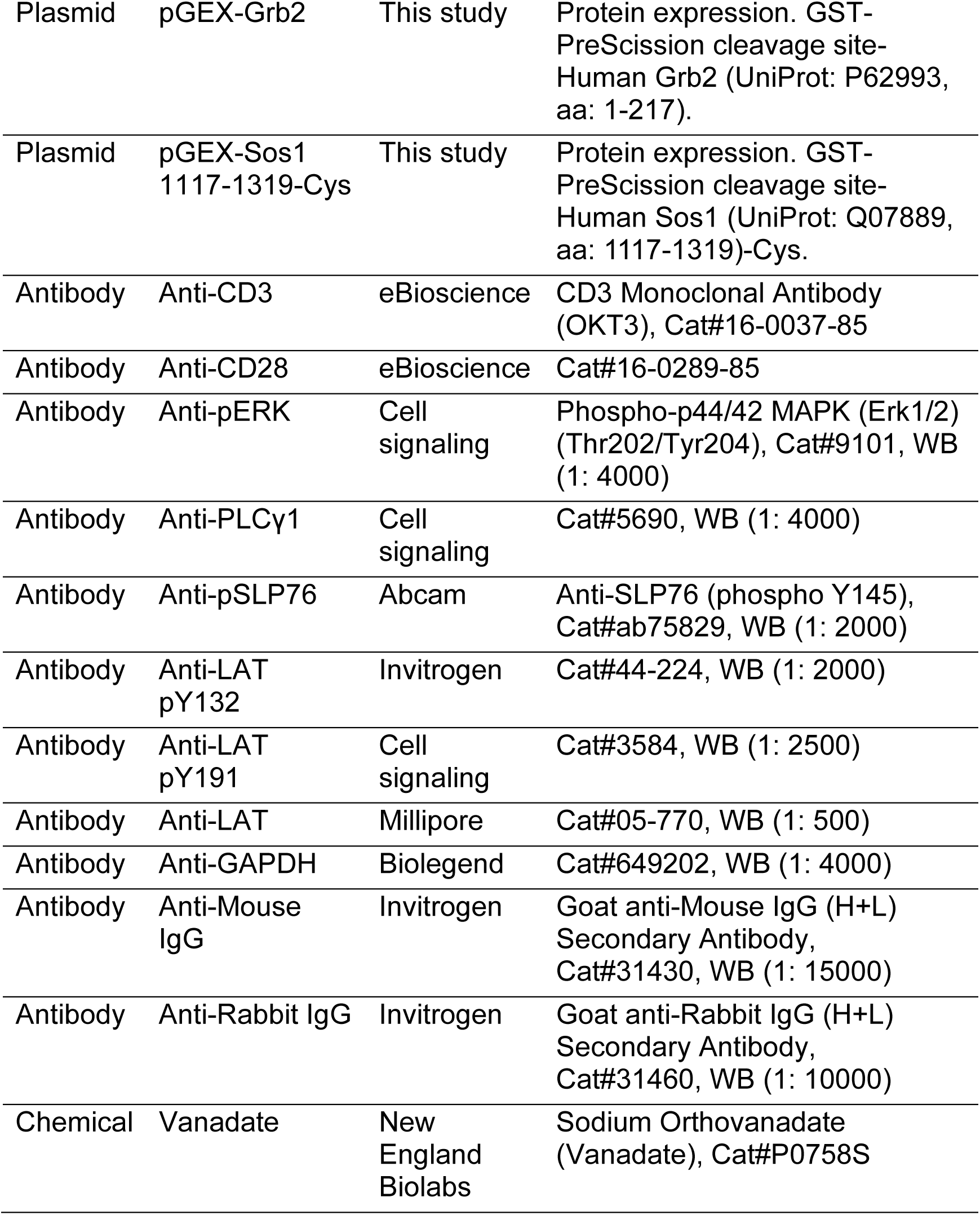

| Reagent type | Designation | Source or reference | Additional information |
| --- | --- | --- | --- |
| Cell line | HEK293T | UCSF CCF |  |
| Cell line | Jurkat T cell E6.1 | UCSF CCF |  |
| Cell line | PLCG1 null (J74B) | This study | PLCG1 knockout by CRISPR |
| Cell line | GRB2 null (J81) | This study | GRB2 knockout by CRISPR |
| Cell line | GADS null (J82) | This study | GADS knockout by CRISPR |
| Plasmid | pMD2.G | Vale Lab | Viral packaging, Plasmid |
| Plasmid | poPAX | Vale Lab | Viral packaging, Plasmid |
| Plasmid | pSpCas9(BB)-PLCG1 234F-2A-GFP (XSB717) | This study | CRISPR-cas9 vector for knocking out PLCG1. |
| Plasmid | pSpCas9(BB)-Grb2 42F-2A-GFP (XSB740) | This study | CRISPR-cas9 vector for knocking out GRB2. |
| Plasmid | pSpCas9(BB)-Gads 97F-2A-GFP | This study | CRISPR-cas9 vector for knocking out GADS. |
| Plasmid | pHR-sfGFP | This study | Lentiviral vector with a super fold GFP tag |
| Plasmid | pHR-PLCγ1-sfGFP | This study | Lentiviral vector. Bovine PLCγ1 (UniProt: P08487, aa: 1-1291) fused with an sfGFP tag. |
| Plasmid | pHR-PLCγ1ΔnSH2-sfGFP | This study | |
| Plasmid | pHR-PLCγ1ΔSH3-sfGFP | This study | |
| Plasmid | pHR-PLCγ1<br>nSH2-cSH2-SH3-sfGFP | This study | |
| Plasmid | pHR-LAT-mCherry | This study | Lentiviral vector. Human LAT (UniProt: O43561-2, aa: 1-233) fused with an mCherry tag. |
| Plasmid | pFastBac-Bv<br>PLCγ1-SNAPf | This study | Protein expression. His10-TEV-Bovine PLCγ1 (UniProt: P08487, aa: 1-1291) fusing SNAPf tag. |
| Plasmid | pGEX-ybbR-BV PLCγ1<br>nSH2-cSH2-SH3 | This study | Protein expression. GST-PreScission cleavage site-ybbR tag-Bovine PLCγ1 (UniProt: P08487, aa: 550-851). |
| Plasmid | pGEX-ybbR-BV PLCγ1<br>nSH2-cSH2 | This study | Protein expression. GST-PreScission cleavage site-ybbR tag-Bovine PLCγ1 (UniProt: P08487, aa: 550-790). |
| Plasmid | pGEX-ybbR-BV PLCγ1<br>nSH2-SH3 | This study | Protein expression. GST-PreScission cleavage site-ybbR tag-Bovine PLCγ1 (UniProt: P08487, aa: 550-667, 757-851). |
| Plasmid | pGEX-ybbR-BV PLCγ1<br>cSH2-SH3 | This study | Protein expression. GST-PreScission cleavage site-ybbR tag-Bovine PLCγ1 (UniProt: P08487, aa: 658-851). |
| Plasmid | pGEX-ybbR-BV PLCγ1<br>nSH2 | This study | Protein expression. GST-PreScission cleavage site-ybbR tag-Bovine PLCγ1 (UniProt: P08487, aa: 550-667). |
| Plasmid | pGEX-ybbR-BV PLCγ1<br>cSH2 | This study | Protein expression. GST-PreScission cleavage site-ybbR tag-Bovine PLCγ1 (UniProt: P08487, aa: 658-790). |
| Plasmid | pET-MBP-His8-LAT 48-233-His6 | This study | Protein expression. MBP-TEV-His8-Human LAT (UniProt: O43561-2, aa: 488-233)-TEV-His6. |
| Plasmid | pFastBac-His10-CD45<br>598-1304-SNAP | This study | Protein expression. His10-TEV-Human CD45 (UniProt: P08575, aa: 598-1304)-SNAP. |
| Plasmid | pGEX-Grb2 | This study | Protein expression. GST-PreScission cleavage site-Human Grb2 (UniProt: P62993, aa: 1-217). |
| Plasmid | pGEX-Sos1 1117-1319-Cys | This study | Protein expression. GST-PreScission cleavage site-Human Sos1 (UniProt: Q07889, aa: 1117-1319)-Cys. |
| Antibody | Anti-CD3 | eBioscience | CD3 Monoclonal Antibody (OKT3), Cat#16-0037-85 |
| Antibody | Anti-CD28 | eBioscience | Cat#16-0289-85 |
| Antibody | Anti-pERK | Cell signaling | Phospho-p44/42 MAPK (Erk1/2) (Thr202/Tyr204), Cat#9101, WB (1: 4000) |
| Antibody | Anti-PLCγ1 | Cell signaling | Cat#5690, WB (1: 4000) |
| Antibody | Anti-pSLP76 | Abcam | Anti-SLP76 (phospho Y145), Cat#ab75829, WB (1: 2000) |
| Antibody | Anti-LAT pY132 | Invitrogen | Cat#44-224, WB (1: 2000) |
| Antibody | Anti-LAT pY191 | Cell signaling | Cat#3584, WB (1: 2500) |
| Antibody | Anti-LAT | Millipore | Cat#05-770, WB (1: 500) |
| Antibody | Anti-GAPDH | Biolegend | Cat#649202, WB (1: 4000) |
| Antibody | Anti-Mouse IgG | Invitrogen | Goat anti-Mouse IgG (H+L) Secondary Antibody, Cat#31430, WB (1: 15000) |
| Antibody | Anti-Rabbit IgG | Invitrogen | Goat anti-Rabbit IgG (H+L) Secondary Antibody, Cat#31460, WB (1: 10000) |
| Chemical | Vanadate | New England Biolabs | Sodium Orthovanadate (Vanadate), Cat#P0758S |

### Recombinant Proteins

A panel of recombinant proteins used in this study was shown in Figure S6B.

The full-length PLCγ1 was purified from insect cells by a baculovirus expression system. PLCγ1 fragments containing individual or combined SH2 and SH3 domains were expressed and purified from bacteria. Peptides including each of the four LAT C-terminal phosphotyrosine residues (pY132, pY171, pY191, pY226) and PLCγ1 pY783 were synthesized by Peptide 2.0 Inc., Chantilly, VA. The peptides are modified with biotin at the N-terminus. The exact protein sequences of PLCγ1 and truncations, and synthesized peptides are listed in Table S1. Gads, Nck1, Grb2, Sos1, CD45, and LAT were purified as previously described (Su et al., 2016).

### Protein purification

#### Full length PLCγ1

The full-length bovine PLCγ1 with an N-terminal His10 tag and a C-terminal SNAP tag was expressed in SF9 cells using the Bac-to-Bac baculovirus expression system (Life Technologies). Cells were harvested by centrifugation and lysed by Dounce homogenizer in 50 mM HEPES (pH 7.4), 300 mM NaCl, 30 mM imidazole, 5% glycerol, 5 μg/mL DNase, 0.5% Triton X-100, 0.5 mM TCEP, 1 mM PMSF, and protease inhibitor cocktail. Centrifuge-cleared lysate was applied to Ni sepharose (GE healthcare), washed with 50 mM HEPES (pH 7.4), 300 mM NaCl, 5% glycerol, 15 mM imidazole, and 0.5 mM TCEP, and eluted with the same buffer with additional 400 mM imidazole. Eluted protein was further purified by size exclusion chromatography using a Superdex 200 prepgrade column (GE Healthcare) in 50 mM HEPES (pH 7.5), 150 mM NaCl, 10% glycerol, and 1 mM TCEP.

#### PLCγ1 nSH2-cSH2-SH3 and purification

The plasmids encoding GST-PLCγ1 nSH2-cSH2-SH3 or truncations were transformed into an *E.coli* BL21 strain. Bacteria were grown at 18 °C in LB medium with 100 μg/mL ampicillin. After cells reached a density of OD_600_ 0.4, protein expression was induced overnight by 0.1 mM IPTG at 18 °C. The culture was harvested and resuspended in PBS buffer, and lysed using a high-pressure homogenizer. The crude lysate was centrifuged at 30,000 rpm for 45 min in a Ti-70 rotor. The supernatant was supplemented with 1 mM DTT and applied to Glutathione Sepharose 4B (GE Healthcare), and washed with 50 mM HEPES (pH 7.4), 150 mM NaCl, 1 mM TCEP and 5% glycerol. The PLCγ1 fragments were cleaved off from the Sepharose by PreScission protease in cleavage buffer (50 mM HEPES, pH 7.4, 150 mM NaCl, 1 mM EDTA, 1 mM DTT and 5% glycerol) at 4 °C overnight. The cleaved products were pooled, concentrated, and further purified by gel filtration using a Superdex 200 size exclusion column (GE Healthcare) equilibrated in 50 mM HEPES (pH 7.4), 150 mM NaCl, 1 mM TCEP and 10% glycerol.

### Protein labeling

The full-length PLCγ1 was labeled with SNAP-Surface Alexa Fluor 647 (NEB, S9136S). Briefly, 10 μM PLCγ1 was mixed with 20 μM SNAP-Ax647 in 500 μL reaction buffer (50 mM HEPES, pH 7.4, 150 mM NaCl, 1 mM TCEP, 1 mM DTT, and 10% glycerol). Following incubation in the dark for 2 hrs at 37 °C, the labeled protein was separated from the solution by size exclusion chromatography using PD MiniTrap G25 (GE Healthcare). PLCγ1 nSH2-cSH2-SH3 and truncations were labeled with CoA 647 (NEB, S9350) via an N-terminal ybbR tag. 10 μM ybbR-fusion proteins were mixed with 20 μM CoA-647 and 2 μM GST-Sfp in 200 μL reaction buffer (50 mM HEPES, pH 7.4, 150 mM NaCl, 1 mM TCEP, 1 mM DTT, 10 mM MgCl_2_ and 10% glycerol). After an incubation of 2 hrs at room temperature, 20 μL Glutathione Sepharose 4B was added to deplete GST-Sfp. The supernatant was further applied to size exclusion chromatography using PD MiniTrap G25 and stored in 50 mM HEPES (pH 7.4), 150 mM NaCl, 1 mM TCEP and 10% glycerol.

### Biochemical reconstitution assay on supported lipid bilayers

The reconstitution of LAT microclusters were carried out following a previous protocol (Su et al., 2017) with minor modification. 1-palmitoyl-2-oleoyl-sn-glycero-3-phosphocholine (POPC), 1,2-dioleoyl-sn-glycero-3-phosphoetha nolamine-N-[methoxy (polyethylene glycol)-5000] (PEG-5000 PE) and 1,2-dioleoyl-sn-glycero-3-[(N-(5-amino-1-carboxypentyl)iminodiacetic acid)succiny l] (DOGS-NTA) were purchased from Avanti. A lipid mix composed of 98% POPC, 2% DOGS-NTA, and 0.1% PEG-5000 was dried with a stable flow of argon gas, followed by a complete drying in a vacuum desiccator. The dried lipids were resuspended in PBS followed by 30 cycles of freezing and thawing to break down multivesicular bodies (MVBs) into small unilamellar vesicles (SUVs). The remaining MVBs were depleted by a centrifugation at 35, 000 rpm for 45 min at 4°C using a TLA120 rotor. SUVs are stored under argon at 4°C.

The clustering assay was performed in a 96-well glass bottom plate (Matriplate MGB096-1-2-LG-L). The plate was washed with 5% Hellmanex Ⅲ (Sigma) overnight and 5 M NaOH for three times, followed by a thorough rinse with ultrapure water and equilibrated with a basic buffer (50 mM HEPES, pH 7.4, 150 mM NaCl and 1 mM TCEP). SUVs were added to each well, incubated for 1 hr at 37°C to form the supported lipid bilayers (SLBs). Excess SUVs were washed out after. The bilayers were blocked with clustering buffer (50 mM HEPES, pH 7.4, 150 mM NaCl, 1 mM TCEP and 1 mg/mL BSA) for 30 min at 37°C. His-tagged LAT-Ax488 was incubated with the SLBs for 2 hrs and unbound LAT was washed out. Adaptor proteins (Grb2, Sos1, PLCγ1, etc) diluted in an oxygen scavenger mix (0.2 mg/mL glucose oxidase, 0.035 mg/mL catalase, 25 mM glucose and 70 mM β - mercaptoethanol in clustering buffer) were added to the well to form LAT clusters. Clusters were imaged by TIRF microscopy.

### Cell culture and cell line construction

Jurkat T cell lines were maintained in RPMI medium 1640 (Gibco #11875-093) supplemented with 10% fetal bovine serum (Invitrogen) and 1% PenStrep-Glutamine in a humidified incubator with 5% CO_2_ at 37°C. CRISPR-cas9 gene editing system was used to generate the PLCγ1 knockout Jurkat T cell line.

Jurkat T cell lines stably expressing a fluorescence reporter was generated by lentiviral transduction. HEK293T cells were transfected with the pHR plasmids encoding the gene of interests, and the viral packaging plasmids pMD2.G and psPAX2 using Genejuice (EMD millipore). 72 hr after plasmid transfection, cell culture media containing viral particles were harvested and added into Jurkat cells for infection in RPMI media overnight. Jurkat cells expressing fluorescent proteins were sorted by FACS to generate a stable and homogenous expression population.

### Live cell imaging of microcluster formation in Jurkat T cells

A 96-well glass-bottom imaging plate was coated with 5 μg/mL anti-CD3ε antibody (OKT3) in PBS overnight at room temperature. Unbound OKT3 was washed out the next morning.

The imaging plate was equilibrated with image medium (RPMI Medium 1640 without phenol red, 20 mM HEPES, pH 7.4). 0.1 million of Jurkat T cells were added to each well and microclusters were imaged by TIRF microscopy.

### Immunoblot analysis

Jurkat T cells were washed with PBS three times, incubated at 37°C for 30 min, and stimulated with 2 μg/mL OKT3 (eBioscience #16-0037-85) and 2 μg/mL anti-CD28 antibody (eBioscience #16-0289-85). The reaction was stopped by directly adding 2x SDS-PAGE loading buffer (Bio-Rad #1610737) containing protease inhibitor cocktail (Roche #11873580001). The lysates were boiled for 10 min at 95°C and clarified by centrifugation at 12, 000 rpm for 15 min at 4°C. The supernatants were loaded onto a 4-20% protein gel (Bio-Rad #4568096) for SDS-PAGE analysis, followed by a transferring onto PVDF membrane (Bio-Rad #1620177). The membrane was blocked with TBST buffer containing 5% nonfat milk for 1 hr at room temperature, and blotted with indicated primary antibodies overnight at 4°C. The next day, the membrane was further blotted with HRP-conjugated secondary antibody for one hour at room temperature. Target proteins were detected with a chemiluminescent HRP substrate (Thermo Scientific #34577) and visualized by a Bio-Rad ChemiDoc imaging system.

### Data analysis

Images were analyzed using FIJI (Image J). LAT clustering was quantified as normalized variance, which equals to the square of standard deviation divided by mean (after subtracting background). Graphs in the same panel were displayed with the same brightness and contrast setting. The half recovery time of FRAP was obtained by fitting a “one-phase association” model in GraphPad Prism 7.00 software.

### Computer simulations

The computational model consists of two-dimensional particles of circular shape and diameter σ, with patches playing the role of binding sites (Figure 4A). All particles interact with each other through Weeks-Chandler-Anderson potential, of strength 10 *k*_B_*T*, representing volume exclusion. On top of that, two particles can attract each other if two interacting patches on their surfaces come close: in this case we say that they form a bond. LAT particles have 4 binding sites, representing pY132, pY171, pY191 and pY226; PLCɣ1 particles have 3 binding sites, representing the nSH2, cSH2 and SH3 domains; Sos1 particles have 4 identical binding sites, representing PRMs; Grb2 particles have 3 binding sites, representing the SH2, nSH3 and cSH3 domains.

2d molecular dynamics simulations are performed using LAMMPS (Plimpton, 1995). There, patches are implemented by means of ghost particles of diameter equal to 0.05 σ and positioned at fixed distance 0.475 σ from the center of the proper particle. Two interacting patches (as per Figure 4A) interact via a cosine-squared potential with range of attractiveness 0.15 σ between the centers of the patches. The maximum depths ɛ of the cosine-squared potential is set to 30 *k*_B_*T*, *k*_B_ being the Boltzmann constant and *T* being the temperature, rendering the bonds practically unbreakable within simulation time. Each molecule, i.e. a volume-excluded particle together with its patches representing binding sites, is treated as a rigid body. The angular distance between patches belonging to the same molecule is chosen in such a way to 1) make binding between two patches exclusive, and 2) allow a Sos1 and a Grb2 molecule to bind through either one or two pairs of patches. 2d Brownian dynamics is ensured by an overdamping Langevin thermostat, enforcing room temperature at every step.

In simulations, the number of LAT molecules is fixed to 200. The ratio of LAT:Sos1:Grb2 particles is kept constant at 1:1:2, whereas the proportion PLCγ1:LAT is varied from 0 to 3. Our goal is not to capture the exact experimental value of these ratios, both because in experiments the surface density of cytosolic cross-linkers at the membrane is unknown and because it can only affect quantitatively, but not qualitatively, the results. Different surface densities ρ_LAT_ are chosen for LAT, from 0.005 to 0.05 *σ*^‒2^, with no noticeable difference in the mechanism of clustering (see Figure S4A and SI). In the text we show results for ρ_LAT_ = 0.02 *σ*^‒2^.

The coalescence likelihood is defined as the number of possible bonds that can form, in principle, between a cluster and another one, summed over all clusters available at a given time. This amounts to

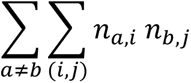

where *n_a,i_* is the total number of yet unbound binding sites of type *i* present in cluster *a*, the first sum runs over all combinations of two clusters, labeled *a* and *b*, and the second sum runs over all types of interacting patch pairs (*i*, *j*), as per Figure 4A. For simplicity, spatial hindrance is neglected in the computation of the coalescence likelihood: all unbound binding sites are counted, irrespective of whether they are physically accessible or not. This assumption is justified by the fact that for clusters of the size and structure that we observe, the vast majority of molecules is situated at the boundary and can therefore contribute to cluster coalescence.

The compactness parameter mentioned in the text is based on the analysis of the gyration tensor of each cluster, using as pole the cluster’s center of mass (the gyration tensor is equal to the inertia tensor for unit masses). The tensor is diagonalized, in order to obtain the three principal moments of inertia: one of them, *I_z_*, represents the inertia of in-plane rotations about the axis *z*, perpendicular to the plane and going through the center of mass; the remaining two principal moments *I_1_* and *I_2_* represent the squares of the gyration radii of the cluster along two perpendicular directions in the plane. It is possible to compute the moment of inertia *I_z_*_,min_ along *z* of an ellipse of gyration radii 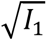 and 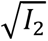, made of close-packed circles (continuous bulk density for hexagonal close-packing is assumed): this represents the inertia of a fictional cluster of approximately the same shape as the real one, made of as many circular particles as there can fit. The compactness parameter is then defined as the ratio *I_z_*_,min_ / I_z_ and tends to 0 for a maximally sparse cluster and to 1 for a maximal-mass-density ellipse with no void regions (see Figure 4B for examples and Section SI 4).

**Table S1.**
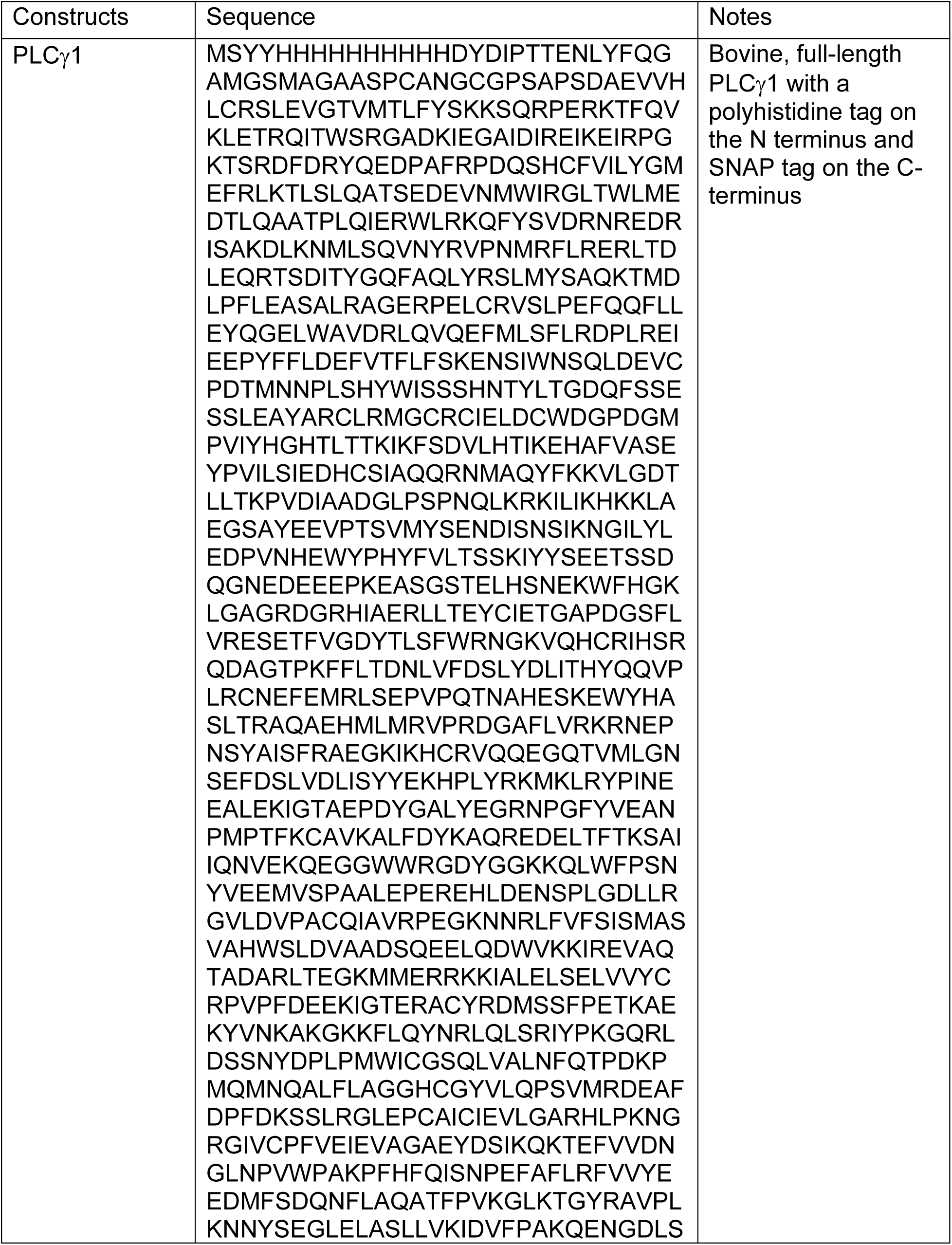
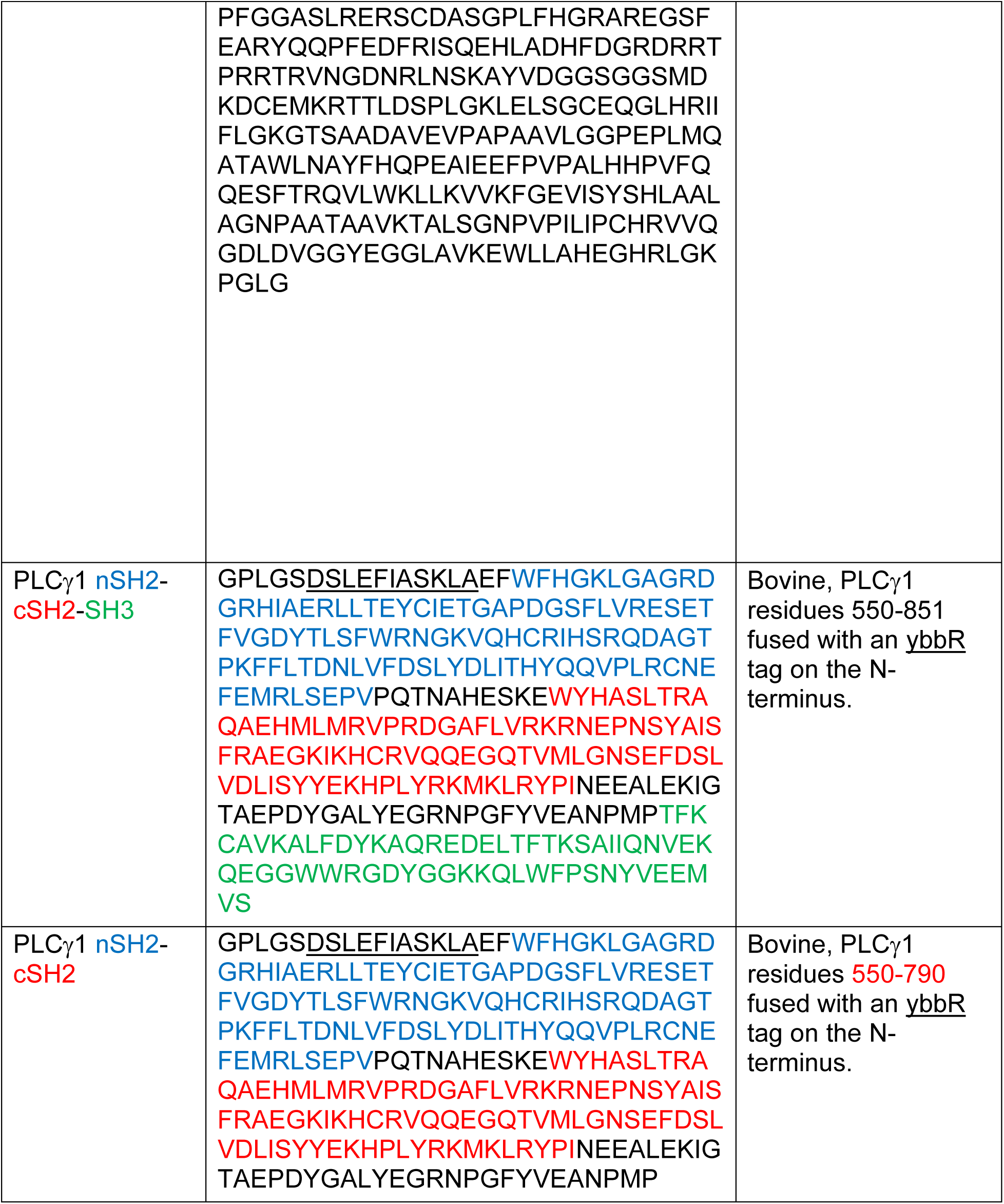
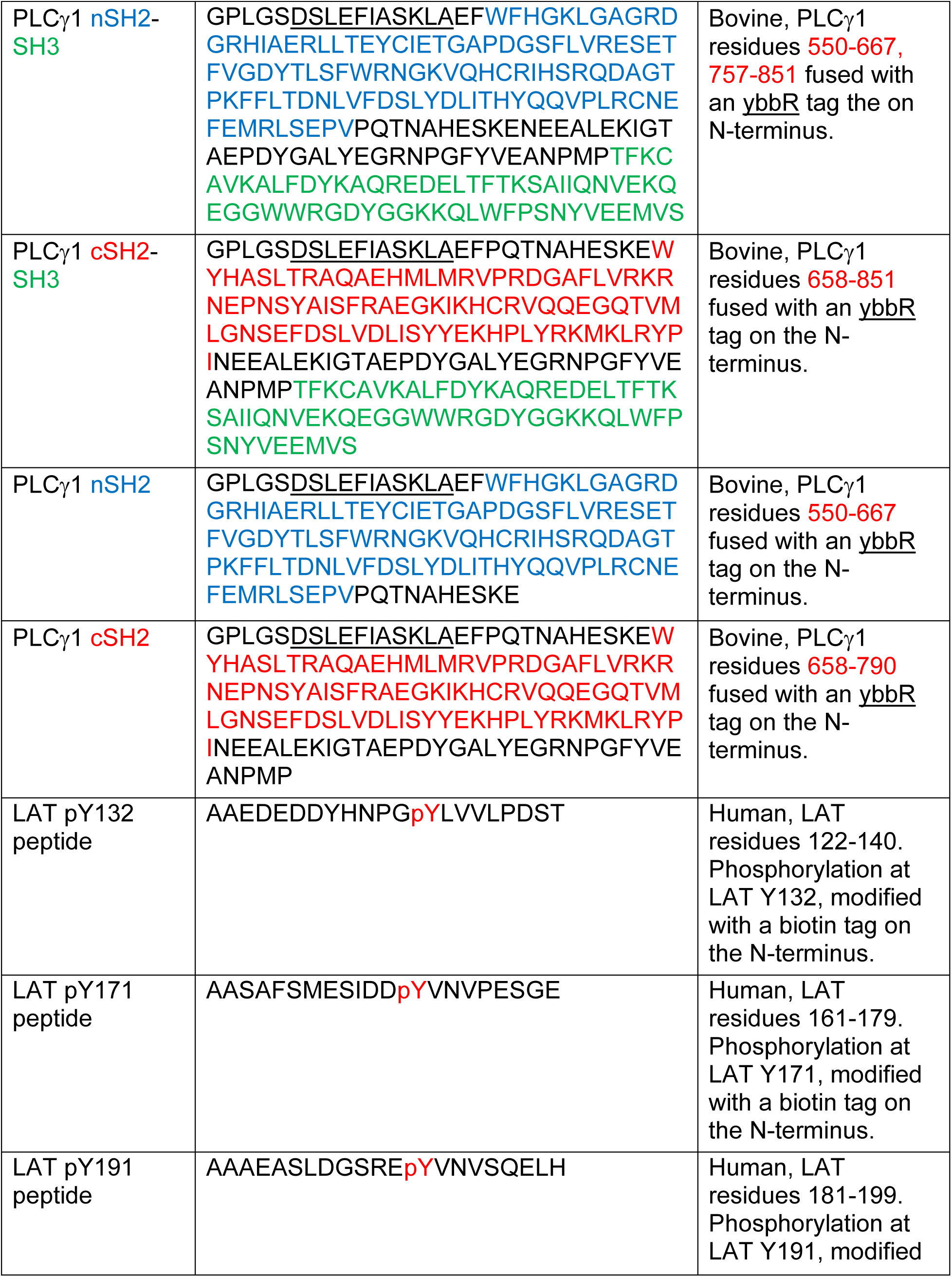
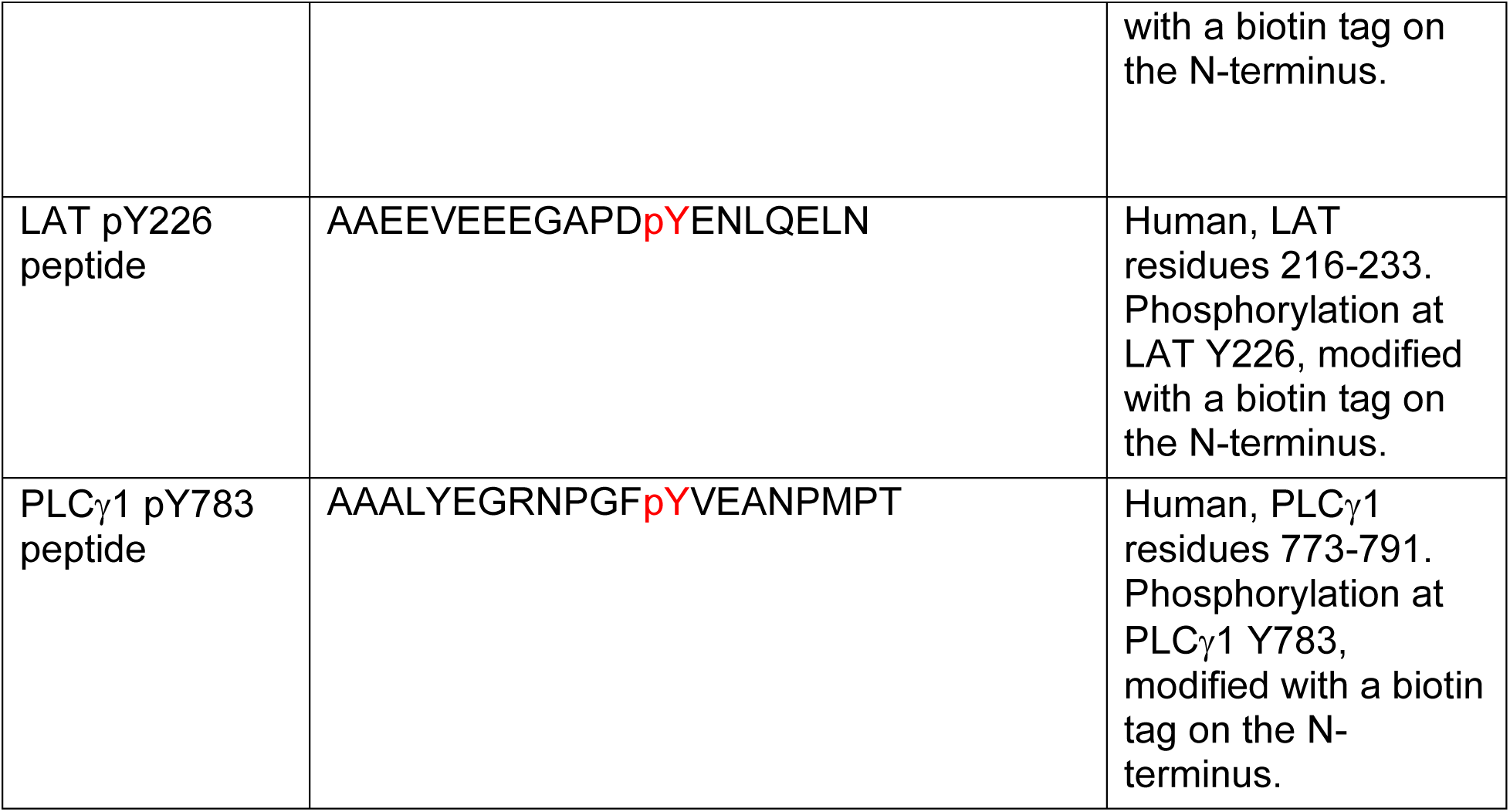
New recombinant proteins and peptides used in the study.

## Supplemental Information

### Supplementary Figure Legends

**Figure S1.**
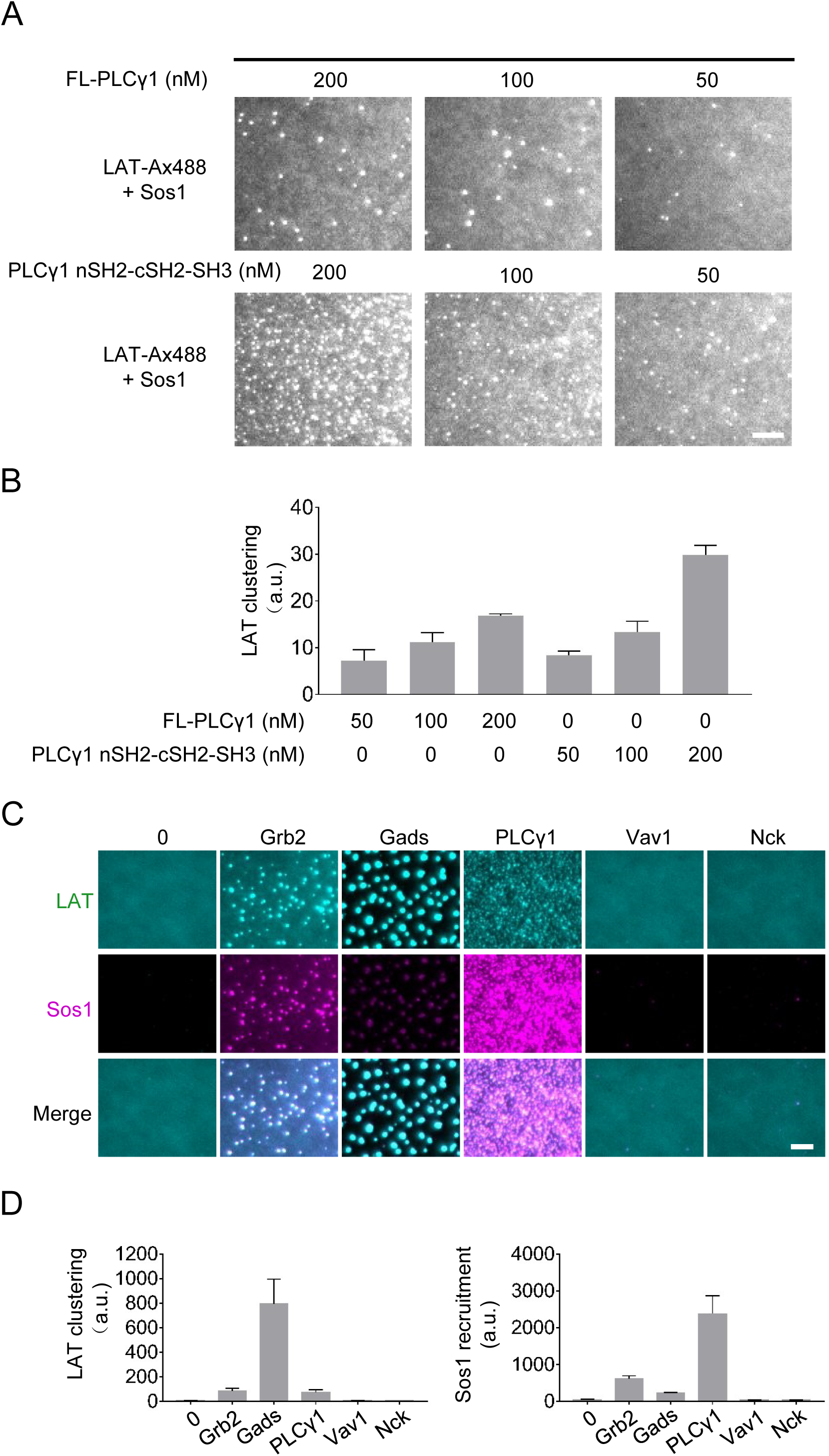
PLCγ1 promotes LAT cluster formation in vitro. (A) TIRF microscopy revealed LAT microcluster formation with the full-length or SH2-SH2-SH3 domain of PLCγ1. Alexa-488 labeled, phosphorylated LAT at 300 molecules / μm^2^ was incubated with 250 nM Sos1 and indicated concentrations of PLCγ1 or fragment. Scale bar: 5 μm. (B) Quantification of PLCγ1 driven-LAT microclusters. Shown are mean ± SD. N= 3 independent experiments. (C) TIRF microscopy revealed LAT microcluster formation with different SH2 and SH3 domain proteins. Alexa-488 LAT at 300 molecules / μm^2^ was incubated with 250 nM Sos1 (labeled with Alexa-647) and indicated proteins at 500 nM. Scale bar: 5 μm. (D) Quantification of LAT clustering and membrane recruitment of Sos1. Shown are mean ± SD. N= 3 independent experiments.

**Figure S2.**
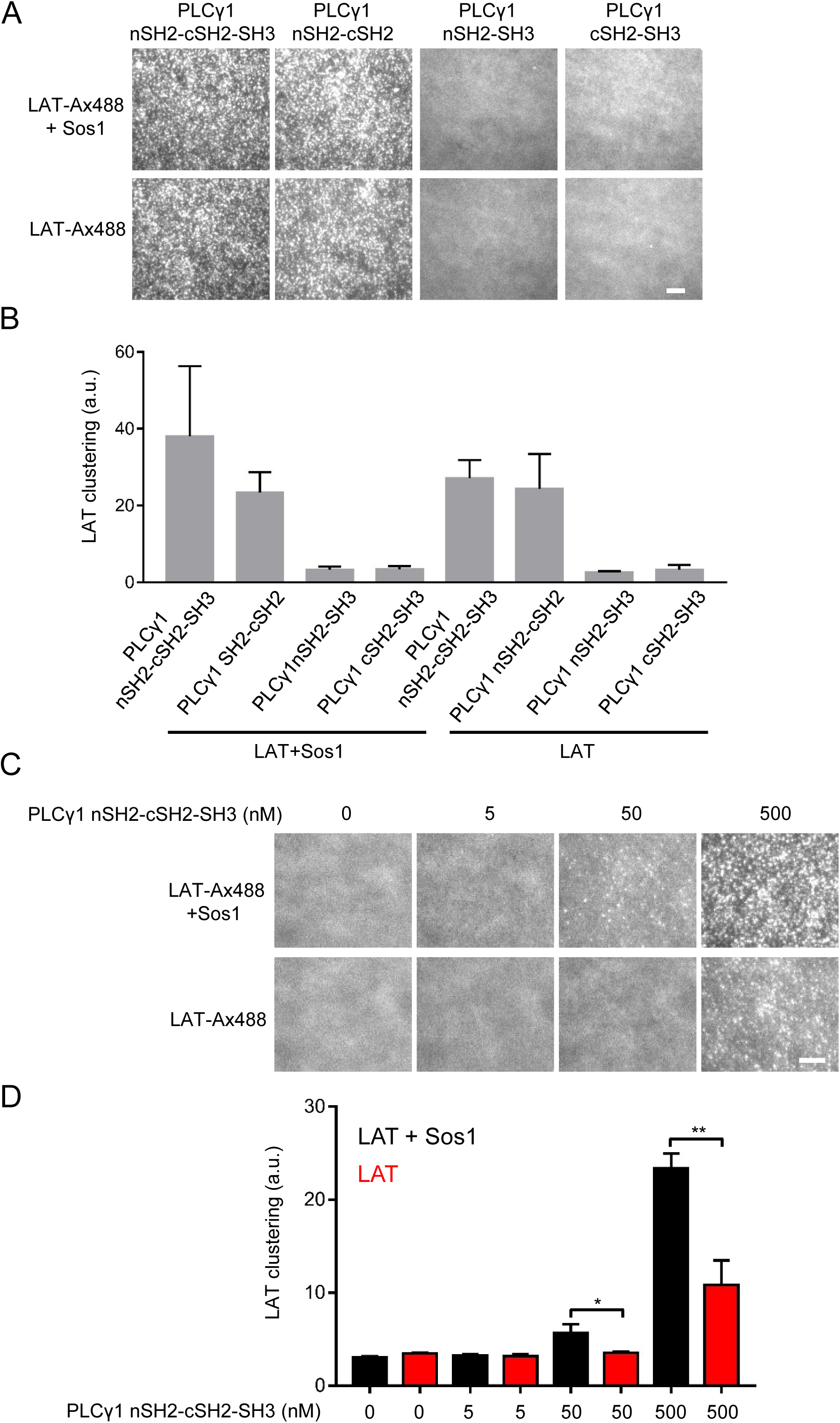
Domains required for PLCγ1-driven LAT clustering. (A) TIRF microscopy revealed LAT microcluster formation with high concentration of PLCγ1 fragments. Alexa-488 LAT at 300 molecules / μm^2^ was incubated with 125 nM Sos1 and 500 nM of indicated PLCγ1 fragments. Scale bar: 5 μm. (B) Quantification of LAT clustering in (A). Shown are mean ± SD. N= 3 independent experiments. (C) TIRF microscopy revealed LAT microcluster formation with titrated PLCγ1. Alexa-488 LAT at 300 molecules / μm^2^ was incubated with or without 250 nM Sos1 and indicated concentration of PLCγ1 nSH2-cSH2-SH3 domains. Scale bar: 5 μm. (D) Quantification of LAT clustering in (C). Shown are mean ± SD. N= 3 independent experiments. Unpaired two-tailed t-test. *: p < 0.05, **: p < 0.01

**Figure S3.**
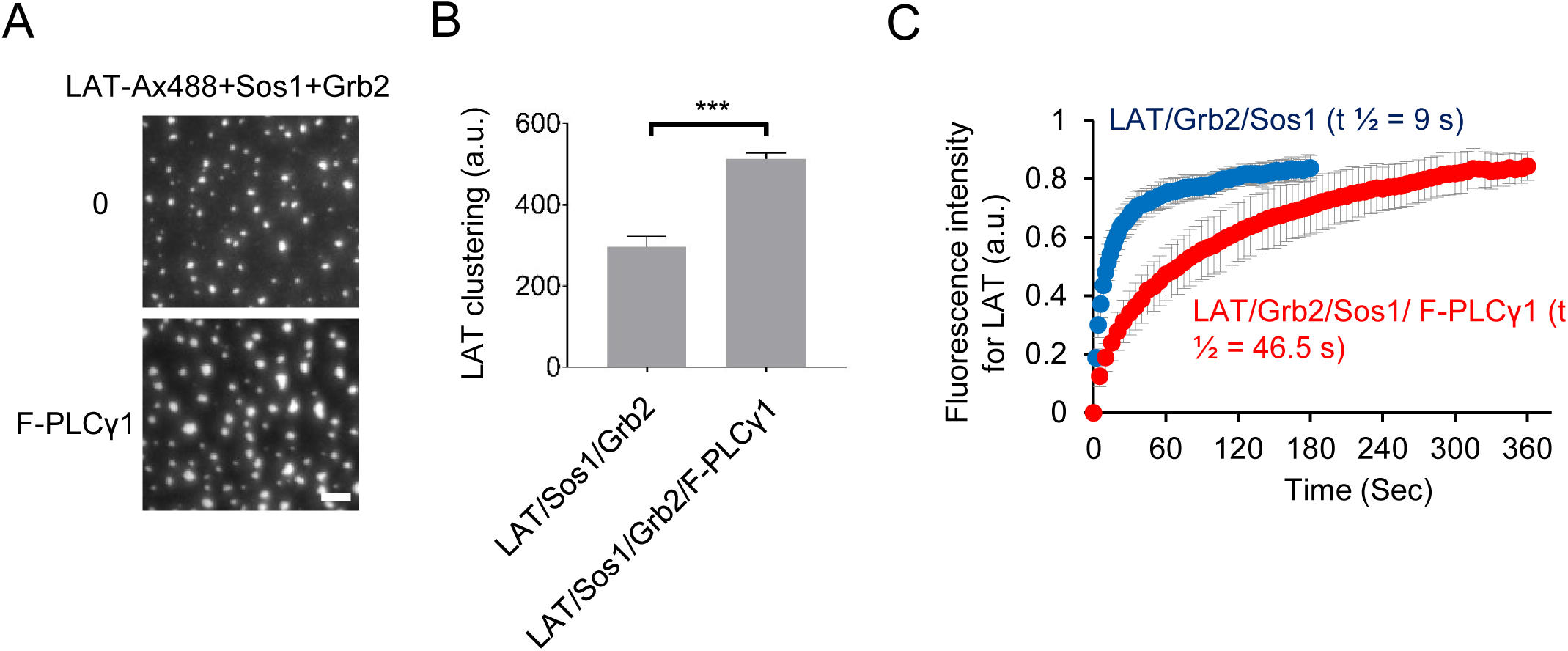
PLCγ1 reduces the mobility of LAT in clusters. (A) TIRF microscopy revealed LAT microcluster formation in the presence or absence of PLCγ1. Alexa-488 LAT at 1000 molecules / μm^2^ was incubated with 500 nM Sos1 and 1000 nM Grb2 with or without 100 nM full-length PLCγ1. Scale bar: 5 μm. (B) Quantification of LAT clustering. Shown are mean ± SD. N= 3 independent experiments. Unpaired two-tailed t-test. ***: p < 0.001. (C) FRAP analysis revealed that PLCγ1 decreases the recovery of LAT signal in clusters after photobleaching. Shown are mean ± SD. N= 10 clusters.

**Figure S4.**
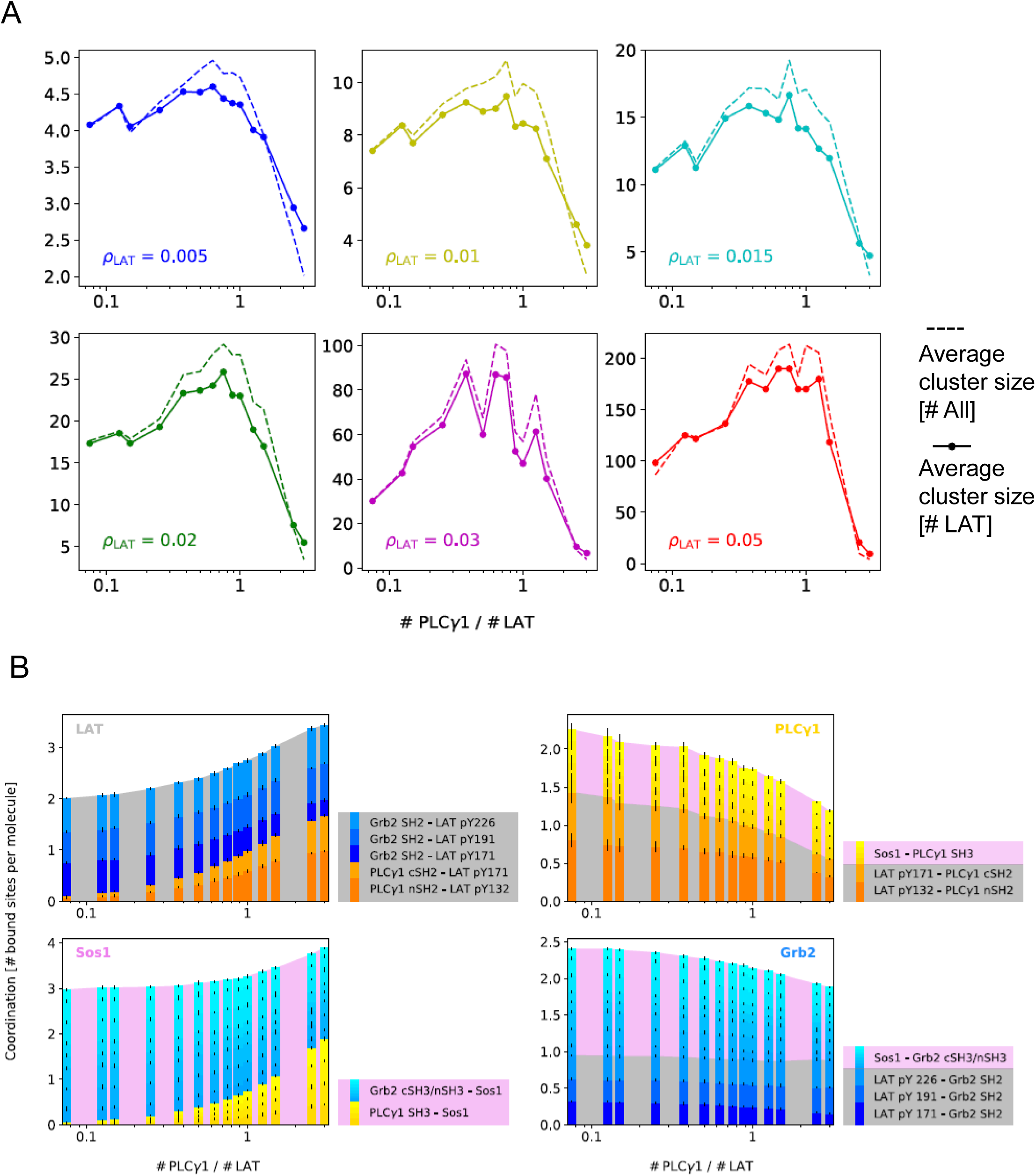
Simulating LAT clustering with PLCγ1. (A) The effect of PLCγ1 on LAT clustering is independent of LAT density ρ_LAT_. A coarse-grained model simulating LAT clustering as a function of PLCγ1 concentration. In a wide range of LAT densities tested, PLCγ1 regulates LAT clustering in a non-monotonic manner. LAT clusters are quantified by the number of LAT in each cluster (solid line) or the total number of molecules (LAT, Grb2, PLCγ1, Sos1) in each cluster (dash line). (B) Average coordination number for all four kinds of particles, as a function of ratio of PLCγ1:LAT, broken down to the contribution of each specific bond. Yellow-orange bars represent bonds involving PLCγ1 and blue bars Grb2; a gray background represents bonds involving LAT, and a pink background Sos1. Here, as throughout Figure 4, ρ_LAT_ = 0.02 σ^−2^. See Section SI 3 for a complete analysis.

**Figure S5.**
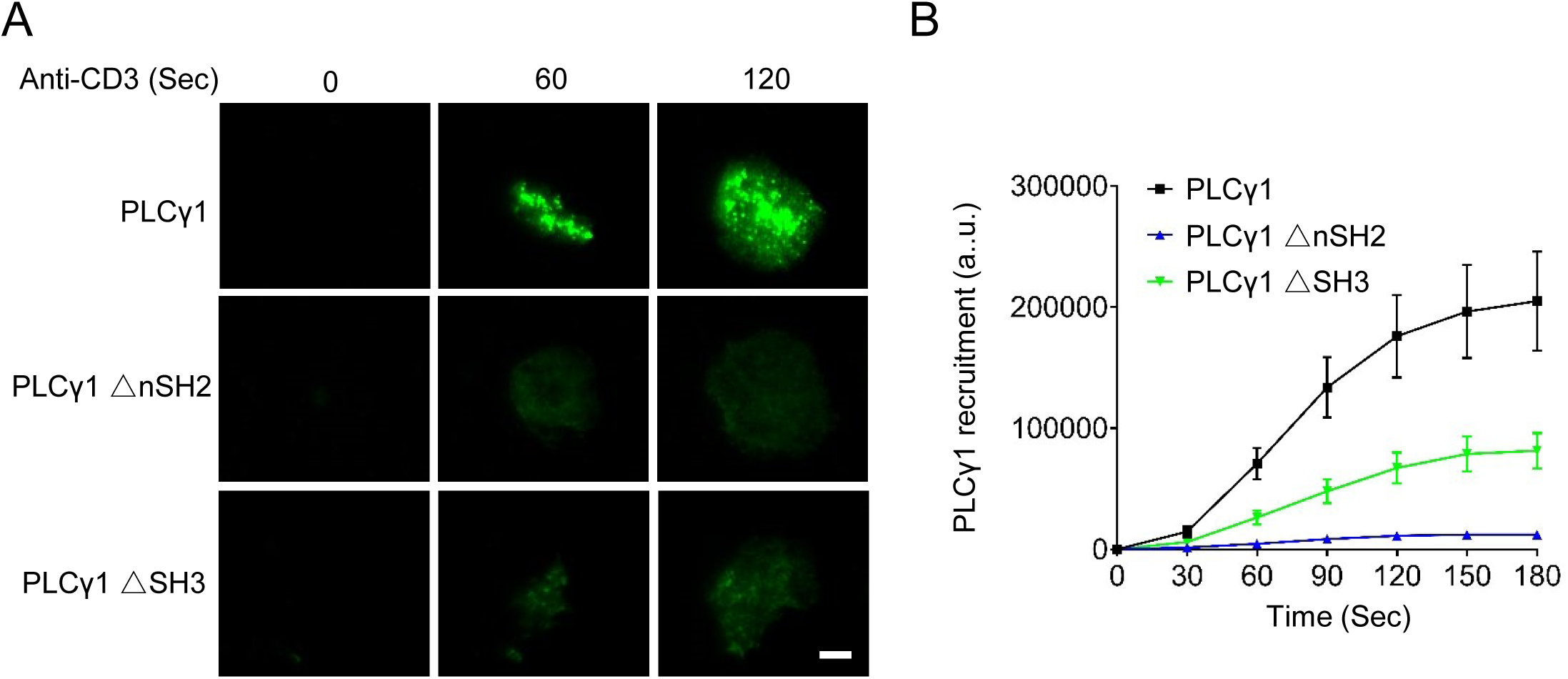
Both the nSH2 and SH3 domain facilitate PLCγ1 recruitment to the T cell plasma membrane. (A) TIRF microscopy revealed recruitment of PLCγ1 to the plasma membrane following TCR activation. Jurkat T cells expressing GFP-tagged full-length, *Δ*nSH2, or *Δ*SH3 PLCγ1 were activated by OKT3-coated cover glass. Scale bar: 5 μm. (B) Quantification of PLCγ1 recruitment to the plasma membrane. Shown are mean ± SEM. N = 24 to 30 cells.

**Figure S6.**
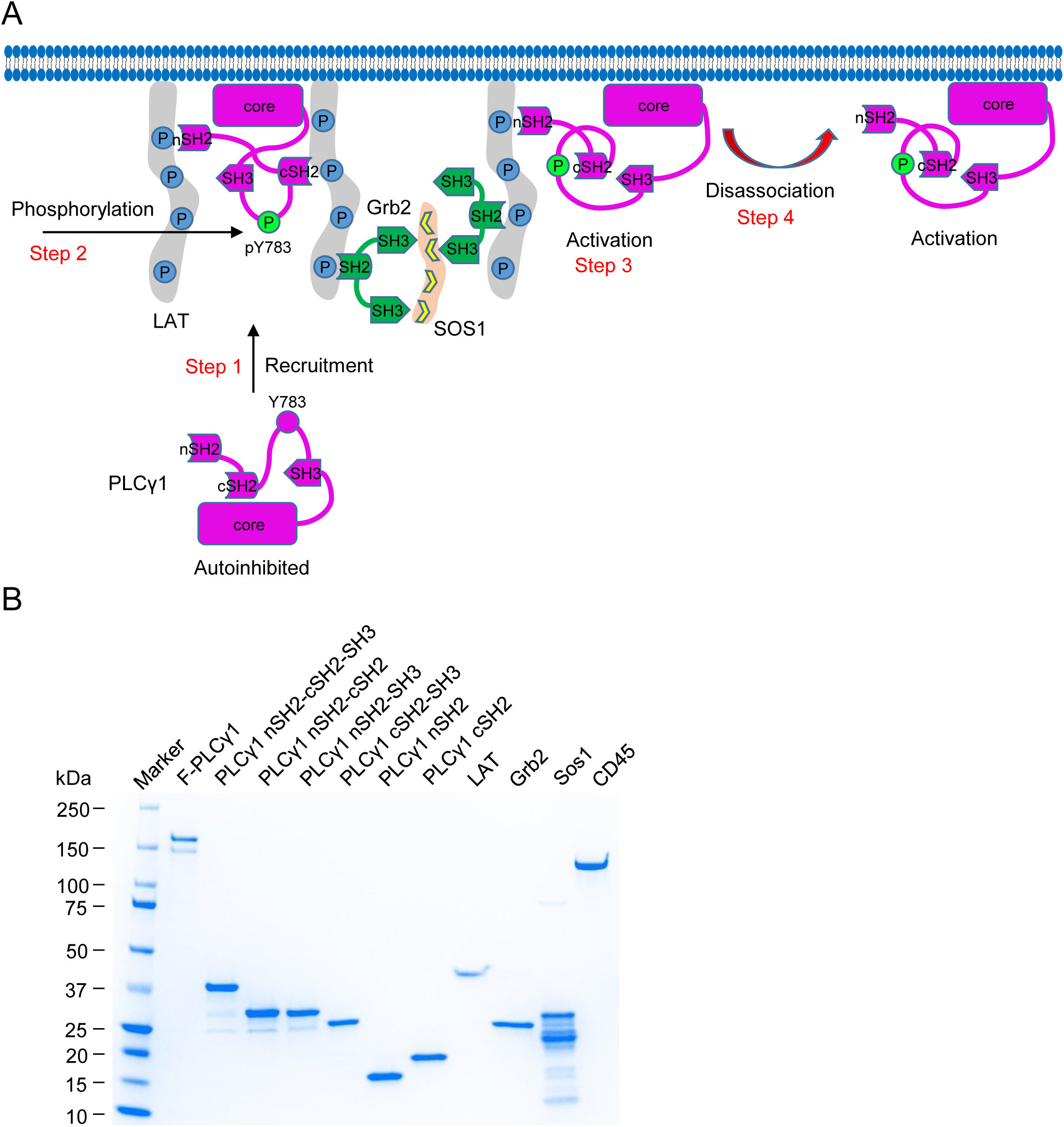
A model for PLCγ1 activity at LAT clusters. (A) Schematics of the recruitment, phosphorylation, activation, and disassociation of PLCγ1 to the LAT clusters. (B) Recombinant proteins used in this study. Purified proteins were applied to SDS-PAGE, followed by Coomassie blue staining.

## Supplementary Movie Legends

**Movie S1**. LAT cluster formation with Grb2 and Sos1. TIRF microscopy revealed LAT cluster formation. His-pLAT-Alexa488 was attached Ni-functionalized supported lipid bilayers at 1000 molecules/μm^2^. Grb2 (1 μM) and Sos1 (0.5 μM) were added at 0 sec to trigger LAT clusters assembly. Shown is a field view of 30 μm × μm.

**Movie S2**. LAT cluster formation with Grb2, Sos1 and PLCγ1. Same condition as in Movie S1 except that PLCγ1 (50 nM) was added together with Grb2 and Sos1 at 0 sec.

**Movie S3**. Early phase of simulation of LAT cluster formation at low PLCγ1 to LAT ratio. The simulation involves 200 LAT, 15 PLCγ1, 400 Grb2, and 200 Sos1 molecules, all in a monomeric state. The simulation starts at second 2.00, corresponding to timestep 0, and ends at timestep 10×10^6^. The interval between two frames is 0.2×10^6^ timesteps and the frame rate is 8 s^-1^. Particles scheme is the same as in Figure 4A: grey-LAT, yellow-PLCγ1, blue-Grb2, and pink-Sos1.

**Movie S4**. Full-length simulation of LAT cluster formation at low PLCγ1 to LAT ratio (high resolution movie for visualizing individual chemical bonds). The simulation involves 200 LAT, 15 PLCγ1, 400 Grb2, and 200 Sos1 molecules, all in a monomeric state. The simulation starts at second 2.00, corresponding to timestep 0, and ends at timestep 50×10^6^. The interval between two frames is 0.2×10^6^ timesteps and the frame rate is 16 s^-1^. Particles scheme is the same as in Figure 4A: grey-LAT, yellow-PLCγ1, blue-Grb2, and pink-Sos1.

**Movie S5**. Early phase of simulation of LAT cluster formation at intermediate PLCγ1 to LAT ratio. The simulation involves 200 LAT, 150 PLCγ1, 400 Grb2, and 200 Sos1 molecules, all in a monomeric state. The simulation starts at second 2.00, corresponding to timestep 0, and ends at timestep 10×10^6^. The interval between two frames is 0.2×10^6^ timesteps and the frame rate is 8 s^-1^. Particles scheme is the same as in Figure 4A: grey-LAT, yellow-PLCγ1, blue-Grb2, and pink-Sos1.

**Movie S6**. Full-length simulation of LAT cluster formation at intermediate PLCγ1 to LAT ratio (high resolution movie for visualizing individual chemical bonds). The simulation involves 200 LAT, 150 PLCγ1, 400 Grb2, and 200 Sos1 molecules, all in a monomeric state. The simulation starts at second 2.00, corresponding to timestep 0, and ends at timestep 50×10^6^. The interval between two frames is 0.2×10^6^ timesteps and the frame rate is 16 s^-1^. Particles scheme is the same as in Figure 4A: grey-LAT, yellow-PLCγ1, blue-Grb2, and pink-Sos1.

**Movie S7**. Early phase of simulation of LAT cluster formation at high PLCγ1 to LAT ratio. The simulation involves 200 LAT, 600 PLCγ1, 400 Grb2, and 200 Sos1 molecules, all in a monomeric state. The simulation starts at second 2.00, corresponding to timestep 0, and ends at timestep 10×10^6^. The interval between two frames is 0.2×10^6^ timesteps and the frame rate is 8 s^-1^. Particles scheme is the same as in Figure 4A: grey-LAT, yellow-PLCγ1, blue-Grb2, and pink-Sos1.

**Movie S8**. Full-length simulation of LAT cluster formation at high PLCγ1 to LAT ratio (high resolution movie for visualizing individual chemical bonds). The simulation involves 200 LAT, 600 PLCγ1, 400 Grb2, and 200 Sos1 molecules, all in a monomeric state. The simulation starts at second 2.00, corresponding to timestep 0, and ends at timestep 50×10^6^. The interval between two frames is 0.2×10^6^ timesteps and the frame rate is 16 s^-1^. Particles scheme is the same as in Figure 4A: grey-LAT, yellow-PLCγ1, blue-Grb2, and pink-Sos1.

## Supplemental Information

### SI 1 Simulation details

#### Setup

Simulations, as described in the Methods section, were performed with a fixed number of LAT molecules (200); the LAT:Sos1:Grb2 relative concentration was fixed to 1:1:2 (i.e. 200 Sos1 molecules and 400 Grb2 molecules) while PLCγ1 concentration was varied (from 0 to 600 molecules). The size of the simulation box was also varied so as to simulate different LAT surface densities *ρ*_LAT_. A rough estimate of the biologically relevant surface density regime stems from the observation that a LAT protein features in reality a 188-aminoacid filament of full-streched length 65 nm. The gyration radius of such filament, approximately proportional to the square root of the number of monomers, is therefore of the order of a few nanometers. If the radius *σ/*2 of a particle in simulations represents the actual gyration radius, an experimental LAT surface density of the order of 10^2^ molecules*/*µm^2^ corresponds to a simulated surface density of the order of 10*^−^*^2^ molecules*/σ*^2^. This is the density regime that we probe.

More computationally costly checks were run on a 4 times larger systems, including 800 LAT molecules, to ensure that the most important features emerging from simulations, such as the re-entrant effect from Fig. 4B were not an artifact of finite size. Finite size effects were not relevant, except for densities and times large enough to allow the formation of big clusters, of size comparable with the total number of particles in the box.

Particles were initialized at time 0 at random positions on a square lattice, spanning the whole simulation box. The dynamics evolves in steps sufficiently small (0.001 Lennard-Jones time units) so as to ensure stability in the integration of the equations of motion. Periodic boundary conditions are imposed at the four box walls.

For each point in the parameters space, defined by *ρ*_LAT_ and by the concentration ratio PLCγ1:LAT, we run 10 (or 5, for times larger than 0.5 × 10^8^ timesteps) different realizations of the dynamics for statistical purposes. Error bars in the shown graphs refer to the standard deviation of the plotted quantity across these different realizations.

#### Interaction potentials

The Weeks-Chandler-Anderson potential used for hard-core repulsion between molecules is the following:

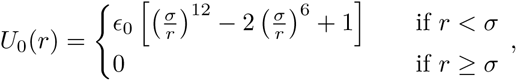

where *r* is the distance between the centers of the two spheres, *σ* is their diameter, and *ϵ*_0_ = 10 *k*_B_*T*.

The cosine-squared potential used for interaction between patches is the following:

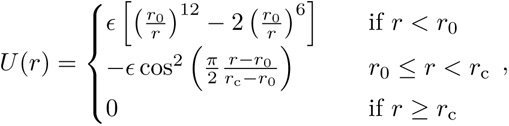

where *r*_0_ = 0.05 *σ*, *r*_c_ = 0.15 *σ* is the range of attraction, and *ϵ* = 30 *k*_B_*T*.

#### Bonds and clusters

A bond between two molecules is defined by two interacting patches being within range of attraction *r*_c_. Two molecules are in the same cluster if and only if they are connected by a path of molecules, such that each molecule forms a bond (as just defined) with the next one. Given the short range and the strength of the interaction, which makes bonds irreversible, clusters are robust against reasonably small variations of the cutoff distance used to define a bond. All the software used in the analysis of simulation results, including clustering, was custom-made.

### SI 2 Kinetics

To study the kinetics of clusters formation, and later the composition of clusters, we compute two quantities: average cluster size and average coordination. The average cluster size is defined by the number of LAT molecules present on average in a cluster at a given simulation time, for a given point in the parameters space. This corresponds to the total number of LAT molecules divided by the total number of clusters, averaged over different realizations. The average coordination, for LAT, PLCγ1, Sos1, or Grb2 molecules, is defined as the number of bonds formed on average by a molecule of that type (i.e. the number of its occupied binding sites). Fig. SI 1 shows the time evolution of these two quantities.

**Figure SI 1:**
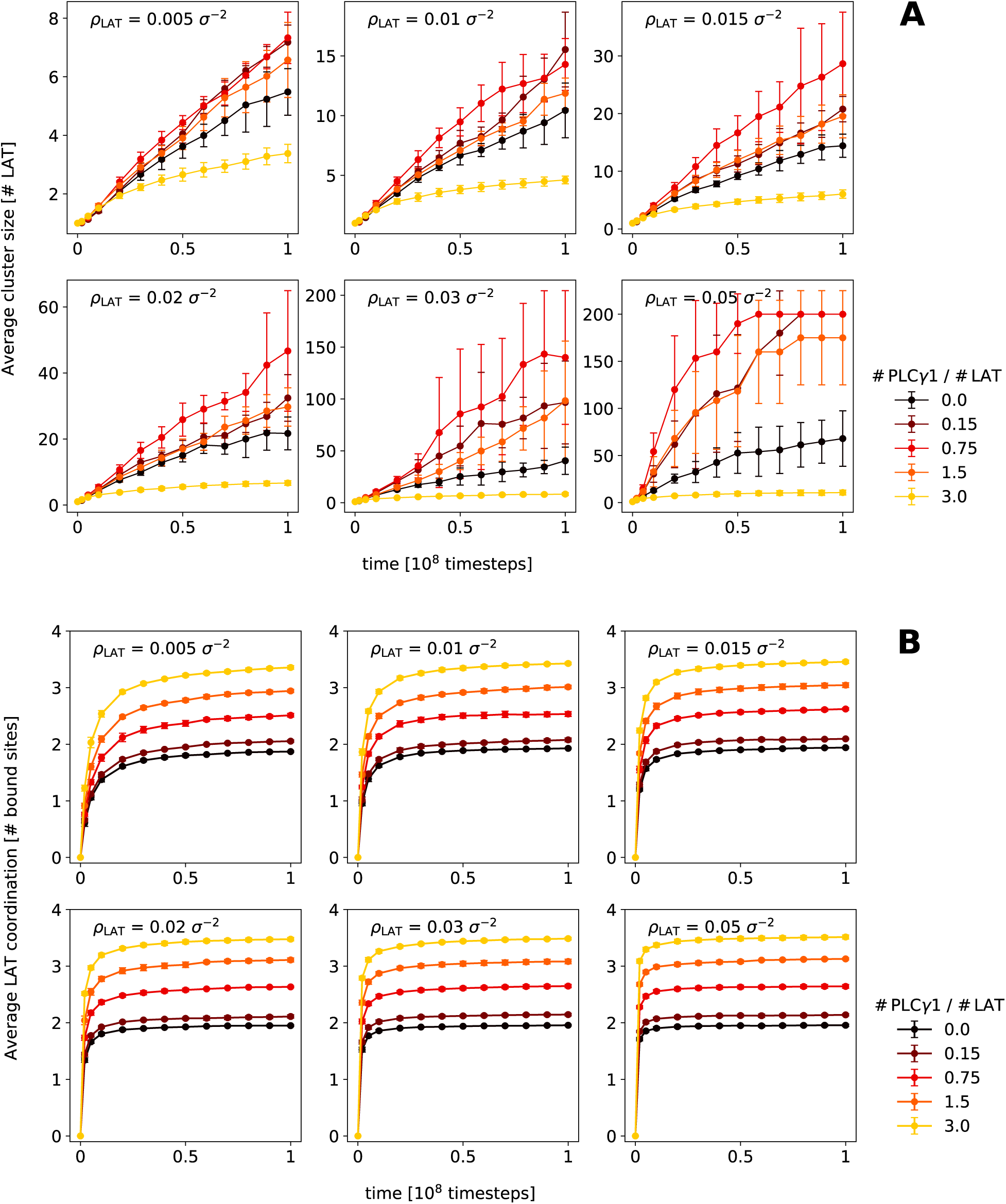
Average cluster size (**A**), measured in number of LAT molecules, and average coordination of LAT molecules (**B**), measured in number of bound sites, as a function of simulation time. Different surface densities and different relative PLCγ1:LAT concentrations are shown. In **A**, at large densities, times and cluster sizes, finite size produces large error bars and a likely spurious saturation effect (see Sec. SI 1).

The average coordination (Fig. SI 1B) exhibits a quick relaxation to its long-time value. This is because most of the bonds are formed during the first 10^7^ timesteps of the simulations. Later on, bonds become rarer, because many binding sites are already occupied, and because the probability of two particles (or clusters) hitting each other by thermal motion decreases as particles condense locally.

On the other hand, the average cluster size (Fig. SI 1A) is dramatically affected by these late-time binding events, as clusters typically double their size upon binding with one another. At low to medium PLCγ1 concentrations, the curves will ideally grow until all particles are condensed into one big cluster: this is a consequence of irreversible interactions and, ultimately, finite size. In a real system, where bonds can break, the equilibrium cluster size will be set by the balance between bonds dynamics (how often bonds break) and density/entropy (how often clusters bump into each other and new bonds form). On the contrary, at high PLCγ1 concentrations, the linkers saturation phenomenon soon forces the system into a locked state, where further clustering is blocked by the unavailability of binding sites.

An interesting feature of Fig. SI 1A is that plots for different *ρ*_LAT_ exhibit the same behavior. Surface density seems to act simply as a rescaling parameter and to influence only the rate of cluster formation: the smaller the density, the less frequently clusters will hit each other risking to coalesce. Analogously, in a real system at equilibrium, the smaller the density, the higher will be the entropic cost associated to bond formation.

Even though our simulations are not meant to reproduce quantitative results, but to give a physically sound explanation to observed experimental phenomena, it can be argued that our model represents realistically an early stage of clustering, where the number of particles is low and bonds have not had the time to break yet. Nonetheless, the non-monotonicity of the average cluster size is a persistent feature, present at all simulated times (except obviously very early stages, where most bonds are not formed yet) and at all densities (Figs. SI 2A and S4A). This suggests that the mechanism behind this re-entrant behavior is robust and at least qualitatively independent from time and *ρ*_LAT_. Motivated by this observation, in the following (as in the main text) we present in detail data captured at time *t*_0_ = 0.5 × 10^8^ steps (when needed, for the sake of conciseness, we also restrict ourselves to a density of *ρ*_LAT_ = 0.02 *σ^−^*^2^). Our conclusions reasonably hold true, irrespective of this particular choice.

**Figure SI 2:**
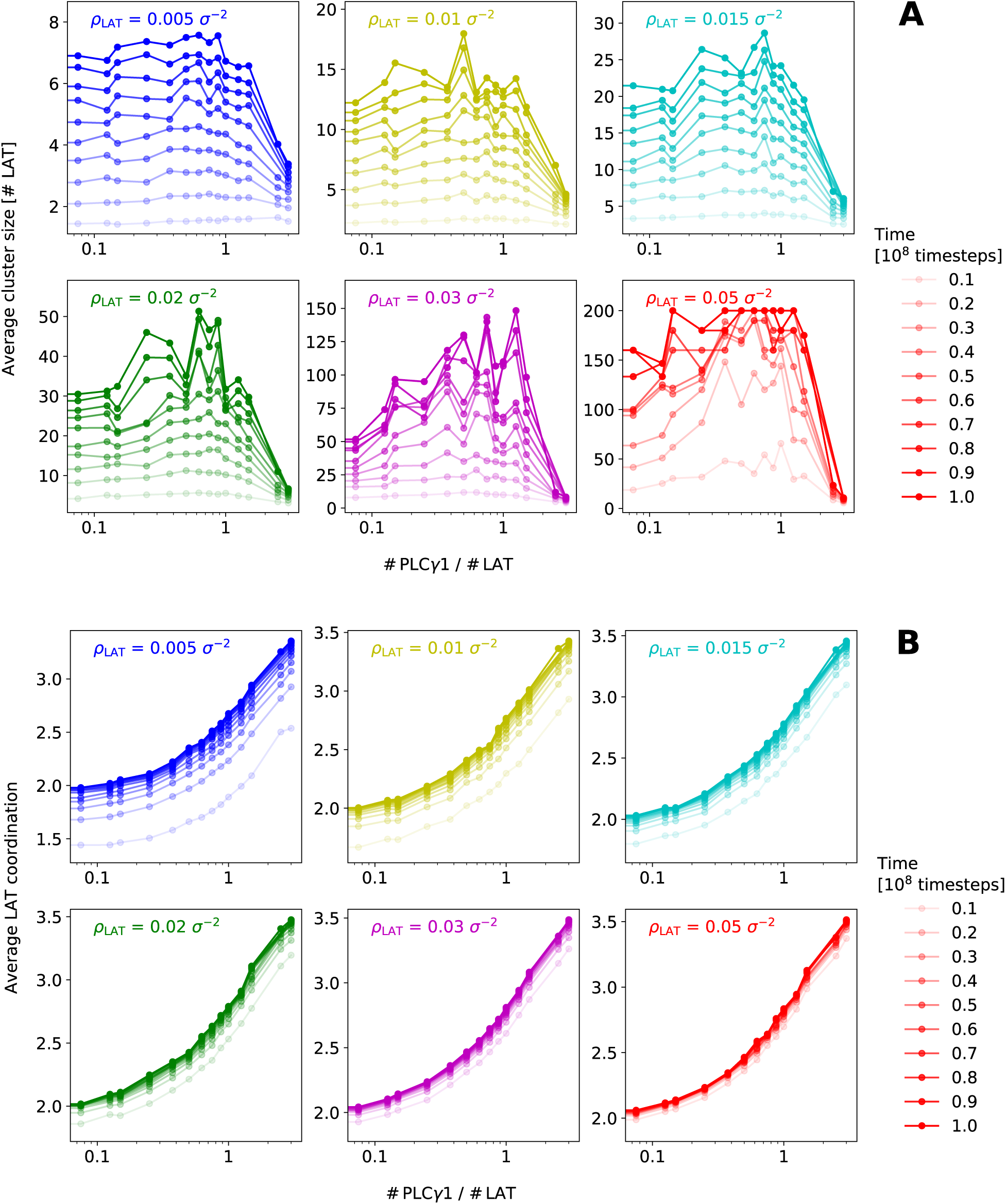
Average cluster size (**A**) and average coordination of LAT molecules (**B**), as a function of relative PLCγ1 concentration, at different times and for different surface densities.

### SI 3 Coordination and coalescence

Fig. SI 3 shows average coordinations of all four kinds of molecules, at fixed time *t*_0_. As previously observed, these curves do not depend on density, except for a slight tendency to decrease as density decreases: this is, again, because at equal time, a lower density system has experienced less collisions than a high density one, and this reflects on the dynamics of the coordination curves (see Fig. SI 2B). To help interpret what happens, we focus on *ρ*_LAT_ = 0.02 *σ^−^*^2^ and break down the four curves from Fig. SI 3, according the the weight of each type of bond. This is done in Fig. S4B, that prompts the following observations.

– The PLCγ1 cSH2 domain compete with the Grb2 SH2 domain for the pY171 site on LAT (top left plot, light orange vs dark blue bar).
– As PLCγ1 concentration increases, LAT’s pY132 and pY171 become completely saturated, due to excess of SH2 binding sites from PLCγ1. Remaining pY191 and pY226 are slightly affected: competition with PLCγ1 for pY171 redirect Grb2’s SH2 to other LAT sites (medium and light blue bars on gray background), so that overall Grb2 binding to LAT stays constant (gray band in bottom right plot).
– At high PLCγ1 concentrations, Sos1 is completely saturated (bottom left plot), due to the overwhelming abundance of SH3 sites from PLCγ1 (yellow bars). In addition, this SH3 domain on PLCγ1 competes for Sos1 with its homologues on Grb2 (light blue bars on pink background), thus causing a decrease in the overall average coordination of Grb2 (bottom right plot).
– Although its presence favors high coordination of LAT and Sos1, and increases in absolute value the number of bonds, PLCγ1’s average coordination decreases with concentration (top right plot). This is due to scarcity of boundable sites on LAT (gray band), and, to a minor extent, of Sos1 (pink band).

**Figure SI 3:**
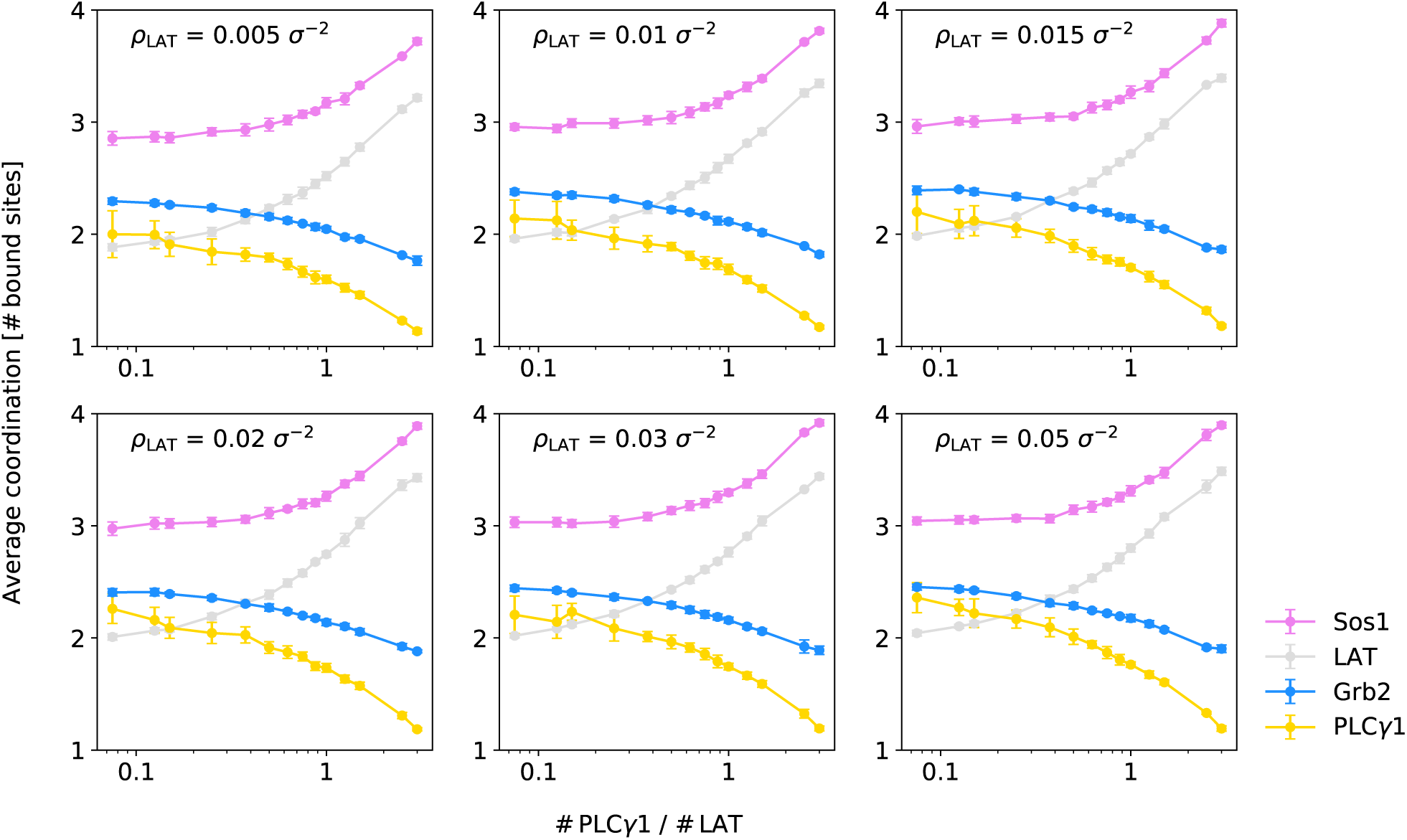
Average coordination number for all four kinds of particles, as a function of relative PLCγ1:LAT concentration at time *t*_0_ = 0.5 × 10^8^ timesteps. Coordination is defined as the number of bound sites, so it ranges from 0 to 4 for LAT and Sos1, and from 0 to 3 for PLCγ1 and Grb2.

All this provides a straightforward interpretation for Fig. 4C, where the coalescence likelihood is observed to decrease drastically, as Sos1 and LAT saturate. The main effect is due to Sos1, which with its 12 binding combinations (Fig. 4A) is involved in the majority of bonds formed in the system.

### SI 4 Shape and compactness

In relation to FRAP experiments, we further analyzed the structure of the networks formed within each cluster, as alluded to in Fig. 4D and in the Methods section. We performed first an analysis of the moments of gyration (or inertia) based on standard Euclidean metric, and then a graph-theoretical analysis based on the connections between molecules rather than on their positions. The latter was motivated by the fact that the geometrical features of our model mainly result from the need to impose bond specificity and exclusivity, and are therefore not necessarily physical. The two analyses are however consistent with each other, showing, as expected, a certain degree of correlation between geometry and network structure.

The mechanical analysis was performed by computing inertia tensors of clusters, relative to their center of mass. To this purpose, since the property of interest was of geometrical and not dynamical origin, all molecules were assigned a unit mass. The center of mass was computed by accounting for periodical boundary conditions during the procedure of clustering analysis. As described in the Methods section, the eigenvalues of the inertia tensor *I_z_*, *I*_1_ and *I*_2_ were determined. *I*_1_ and *I*_2_, the squares of the two in-plane gyration radii, were used to quantify two-dimensional shape through a parameter that we called Roundness and defined as follows:

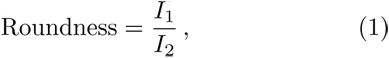

with *I*_1_ *< I*_2_. Taking this ratio amounts to approximating each cluster with an ellipse and estimating its eccentricity. The Roundness tends to 0 for clusters in the shape of a straight line and to 1 for in-plane-rotation symmetric clusters, such as circular ones. We then compared the in-plane-rotation principal moment of inertia (*I_z_*) to the same quantity (*I_z,_*_min_) for a homogeneous ellipse of equal roundness, with a density equal to the bulk density of close-packed circles. The latter is the maximum possible density and gives rise to the minimum possible inertia. In practice, for a given cluster,

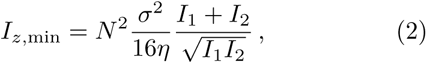

where 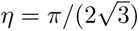 is the packing fraction of a hexagonal close-packed lattice of circles and *N* is the number of molecules in the cluster. This defines the following Compactness parameter:

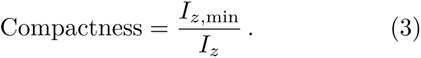

This Compactness intuitively quantifies the presence of holes in the cluster structure: it is 1 if particles are maximally packed (this is impossible in our case, due to the geometry of bonds, so that the achievable Compactness limit is actually below 1), whereas it tends to 0 for a sparse cluster, with many void regions.

Roundness and Compactness, averaged over clusters mixed up from all different realizations, are shown in Fig. SI 4. Once again, *ρ*_LAT_ does not seem to play a role. An effect of PLCγ1 concentration on the Roundness is not recognizable: this would require a symmetry breaking phenomenon that does not seem plausible. On the other hand, Compactness increases with PLCγ1 concentration, as observed in the main text.

**Figure SI 4:**
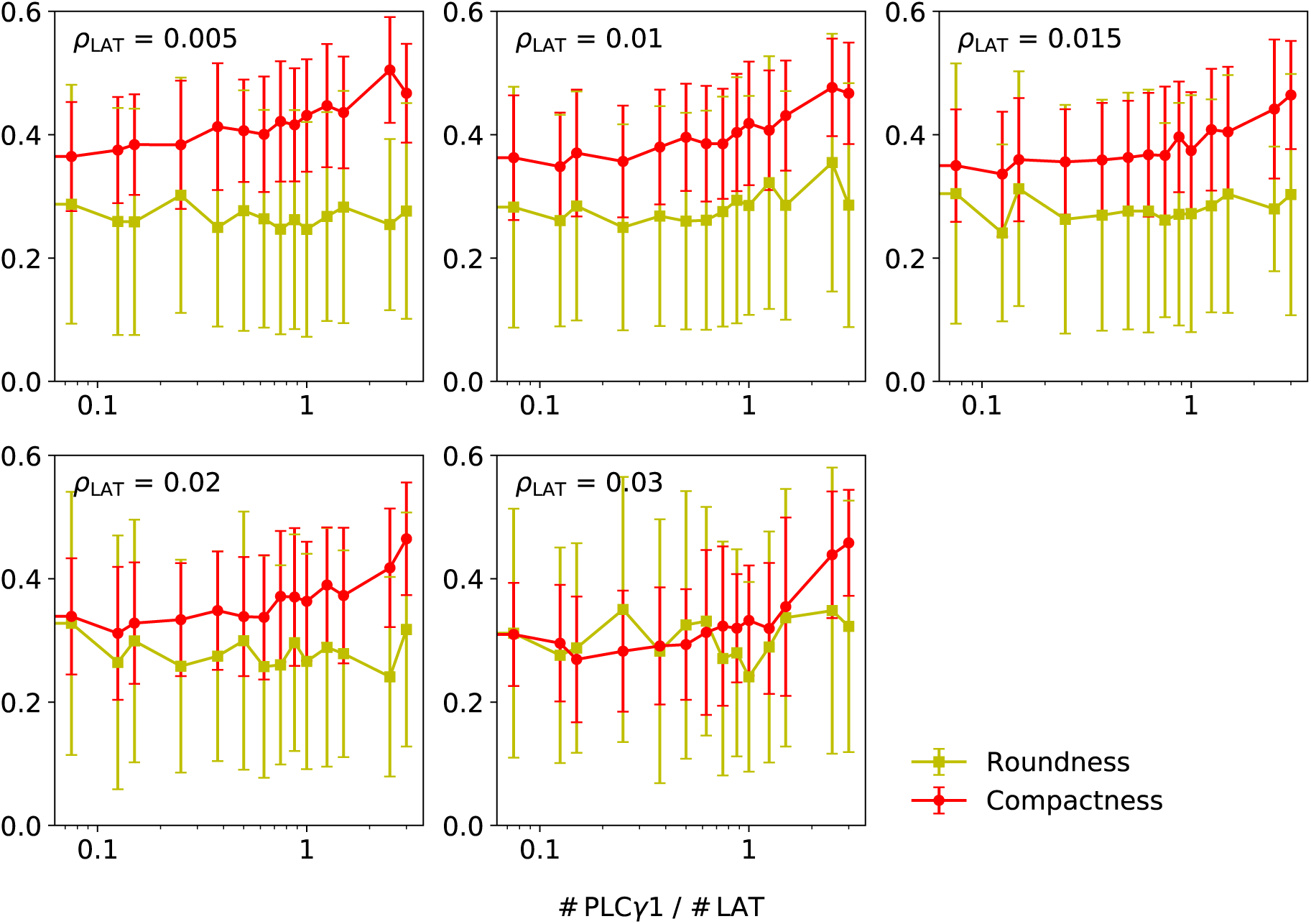
Average Roundness and average Compactness as a function of PLCγ1 concentration, for different surface densities. Here error bars represent the standard deviation across all clusters mixed up from different realizations. Only clusters bigger than 30 molecules are considered, to avoid undesired small-size effects.

The graph-theoretical analysis was performed by defining for each cluster an equivalent undirected unweighted graph, where nodes correspond to particles and edges connect two nodes whose corresponding particles are linked by at least one bond. We characterized the sparsity of such graphs by computing the ratio between number of edges *|E|* and number of nodes *|V|* = *N*. Indeed, a sparse graph will exhibit a linear proportionality between these two quantities, whereas for a fully connected graph *|E| ∼ |V|* ^2^. Surprisingly, although this ratio cannot exceed 2 in our system because of limited particle valence, it does not show any appreciable increase with PLCγ1 concentration. This is in apparent contradiction with the fact that clusters appear more compact as PLCγ1 is added. The reason is that, due to saturation, most of the added PLCγ1 molecules form only one bond, as confirmed by Figs. 4E and S4B, thus increasing the number of nodes with just one edge (terminal nodes or ‘leaves’). This effect is compensated by an increase in the coordination of LAT and Sos1 (see again Figs. 4E and S4B). The result is a constant sparsity. This observation is coherent with and reinforces our picture of available-bonds-limited cluster growth, symbolized by a decreasing coalescence likelihood.

Finally, we attempted to provide a graph-theoretical equivalent to the Compactness parameter defined earlier on a mechanical basis. We defined a graph-theoretical moment of inertia about the center of mass (*I_z,_*_GT_): this exploits on the one hand the concept of graph-theoretical distance *d*(*u, v*), i.e. the number of edges forming the shortest possible continuous path between nodes *u* and *v*, and on the other hand the fact that the moment of inertia relative to the center of mass is the smallest one possible (a corollary of the Huygens–Steiner theorem). The graph-theoretical moment of inertia is given by

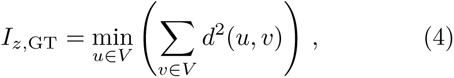

where *V* is the set of nodes. This needs to be compared to a similar quantity for a compact cluster; since the latter will scale as |*V|*^2^, for simplicity we define the graph-theoretical compactness as

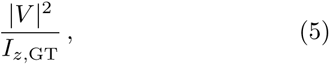

to be compared to Eq. (3). The average of this quantity is plotted in Fig. SI 5 (blue) and increases with PLCγ1 concentration, as expected.

**Figure SI 5:**
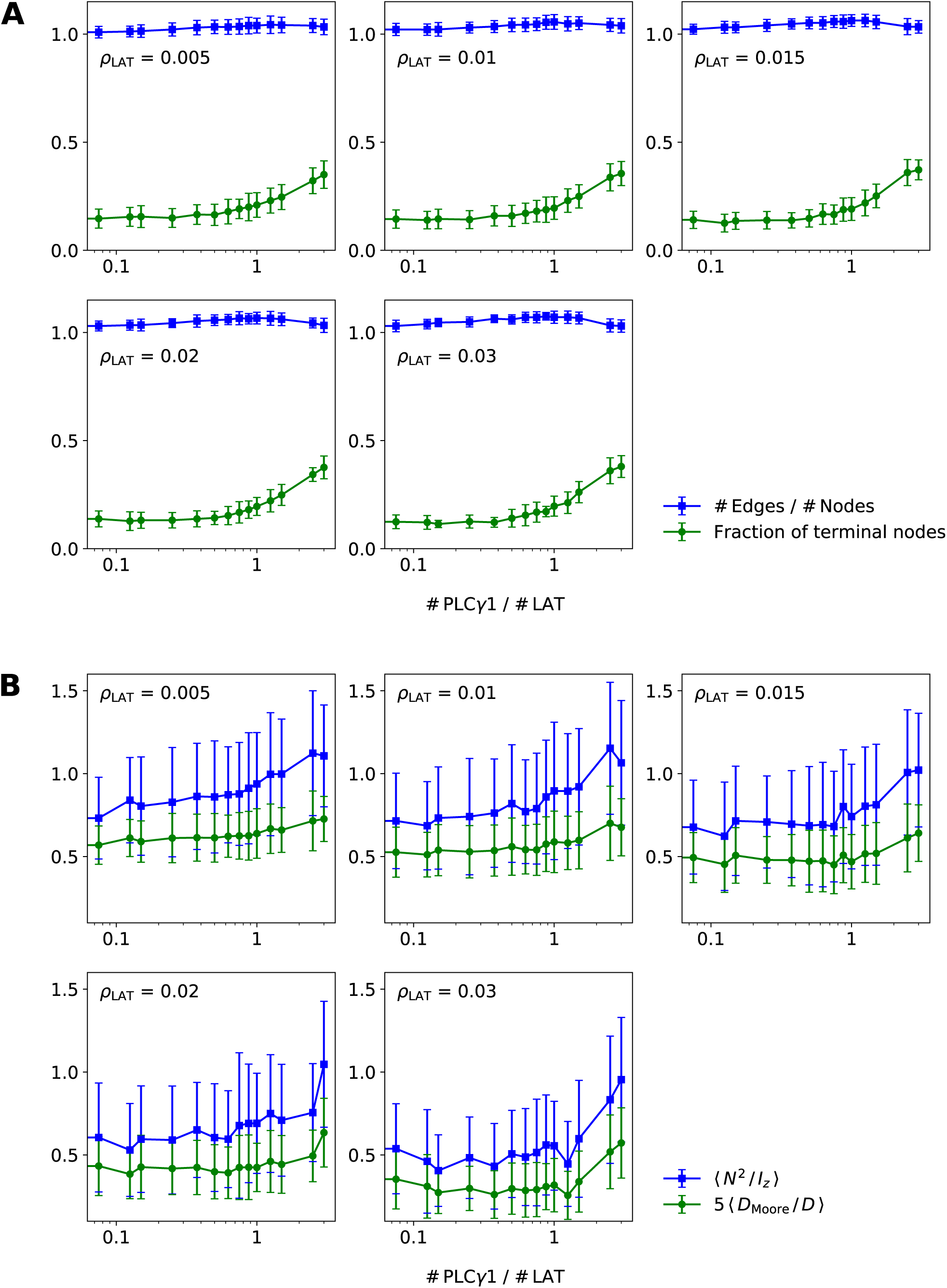
**A**. Average sparsity measurement and fraction of terminal nodes as a function of PLCγ1 concentration. **B**. Graph-theoretical characterization of the cluster compactness, through graph-theoretical moment of inertia and diameter, as a function of PLCγ1 concentration, for different surface densities. Error bars and cluster sample as in Fig. SI 4.

Another measure of the compactness of our graphs originates from the comparison between the diameter *D* of a graph (the distance between the two most distant nodes) and its theoretical lower bound, approximately given by the Moore limit for vertices of maximum degree 4:

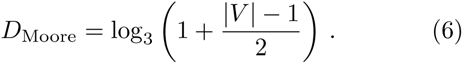

The average of the quantity

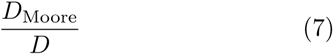

is shown in green in Fig. SI 5 and qualitatively agrees with both the approaches based on graph-theoretical inertia and on mechanical inertia.

